# Single-cell transcriptome landscape of developing fetal gonads defines somatic cell lineage specification in humans

**DOI:** 10.1101/2023.08.07.552336

**Authors:** A. Lardenois, A. Suglia, CL. Moore, B. Evrard, L. Noël, P. Rivaud, A. Besson, M. Toupin, S. Léonard, L. Lesné, I. Coiffec, S. Nef, V. Lavoué, O. Collin, A. Chédotal, S. Mazaud-Guittot, F. Chalmel, AD. Rolland

## Abstract

Gonad development is an exciting model to study cell fate commitment. However, the specification and differentiation of somatic cell lineages within the testis and the ovary are incompletely characterized, especially in humans. In fact, a better understanding of sex determination first requires the identification of all the cell types involved and of their dynamic expression programs. Here we present a comprehensive analysis of approximately 128,000 single cells collected from 33 fetal testes and ovaries between 5 and 12 postconceptional weeks. In particular, a focused analysis of somatic cells allowed us to identify a common population of bipotential progenitors derived from the coelomic epithelium of both male and female gonads and capable of committing to either a steroidogenic or a supporting fate. Moreover, we have shown that early supporting cells, prior to further differentiation into Sertoli or granulosa cells, also give rise to the rete testis/ovarii lineage. Finally, we found that the ovary retains the capacity to feed the supporting cell pool for an extended period of time, directly from the surface epithelial cells and, bypassing the bipotential progenitor step. Altogether, our results provide an unprecedented revisiting of the human gonadal sex determination process.

## Introduction

Gonad development is a unique and remarkable differentiation process that starts from a common bipotential primordium and leads to the formation of two functionally distinct but complementary reproductive organs, the testis and the ovary. In therian mammals this process is genetically controlled: the Y chromosome of male individuals carries the sex determining region Y (SRY) gene which encodes for the so-called testis-determining factor (Berta *et al*, 1990; Koopman *et al*, 1991). Genital ridges first arise from the thickening and differentiation of the coelomic epithelium at the surface of the mesonephroi at around the fourth postconceptional week (PCW) in humans, a time when primordial germ cells (PGCs) also colonize the nascent gonads (Byskov, 1986; Møllgård *et al*, 2010; Gomes Fernandes *et al*, 2018). At this stage, gonads contain multipotent progenitors of supporting and steroidogenic lineages and are considered to be bipotential until 5.5 PCW, when the expression of SRY begins (Hanley *et al*, 2000). Upon expression in supporting cell precursors, SRY induces SOX9 which in turn triggers the differentiation of Sertoli cells, the formation of testis cords and the development of Leydig cells (Vidal *et al*, 2001; Sekido & Lovell-Badge, 2008; Li *et al*, 2014; Rahmoun *et al*, 2017). In the absence of SRY in the XX gonad, the coordination of the RSPO1/WNT4/β-catenin pathway and FOXL2 promotes ovarian development, including the differentiation of supporting precursors into pre-granulosa cells and the subsequent commitment of germ cells into meiosis (Vainio *et al*, 1999; Schmidt *et al*, 2004; Uda *et al*, 2004; Ottolenghi *et al*, 2007; Chassot *et al*, 2008; Liu *et al*, 2009; Le Bouffant *et al*, 2010; Childs *et al*, 2011). It is also well established that male and female molecular expression programs antagonize each other, thereby enabling to reinforce the initiated fate and repressing the alternative one (Kim & Capel, 2006; Chang *et al*, 2008; Maatouk *et al*, 2008; Wilhelm *et al*, 2009; Kashimada *et al*, 2011; Jameson *et al*, 2012; Greenfield, 2015; Bagheri-Fam *et al*, 2017). Altogether, the decisive commitment to a male or female fate during the gonadal sex determination process directs the differentiation of several cell lineages towards one of these two developmental pathways (for review, (Lin & Capel, 2015)).

While progress has been made in the identification of factors involved in gonad formation and differentiation, our understanding of sex determination remains incomplete. Accordingly, the etiology of disorders of sex development remains unexplained in more than half of the cases (for review see, (Reyes *et al*, 2023)). One of the primary reasons for this is a lack of knowledge regarding the different cells that actually make up the gonads. Notably, the lack of specific markers for progenitor cell populations and the inability to perform lineage tracing experiments have prevented the study of gonadal cell specification in humans. Ignoring the existence of certain cell types or subtypes may oversimplify or, even worse, distort our view of gonadal sex determination. The deconvolution capability of single-cell RNA-sequencing (scRNA-seq) has been used in several studies to characterize the large number of cell populations that differentiate within human gonads (Chitiashvili *et al*, 2020; Guo *et al*, 2021; Taelman *et al*, 2022; Wang *et al*, 2022; Garcia-Alonso *et al*, 2022). In addition to the identification of new marker genes for distinct cell lineages, such approaches notably allowed to highlight candidate pathways involved in gonadal cell-cell communication networks, to establish the molecular profiles of fetal rete testis and ovarii cells, or to rule out the mechanisms of X-chromosome dosage in female germ cells. However, these studies included few, if any, samples younger than 7 PCW, and in most cases cells from XX and XY gonads were analyzed separately. In contrast to mouse gonad development, which has now been thoroughly characterized at the single-cell level (Mayère *et al*, 2021a, 2021b; Ademi *et al*, 2022; Niu & Spradling, 2020) (for review, (Estermann *et al*, 2020a)), the early events of gonad formation and the cell fate decisions made in male versus female somatic lineages therefore remain to be elucidated.

Here, we present an in-depth characterization of the transcriptome of human first-trimester gonads from the 5th PCW onwards at the single-cell level. Taking advantage of the large number of early gonads and of the time series design of these scRNA-seq data, we investigated how distinct gonadal cell populations emerge and differentiate before, during and after sex determination. This single-cell transcriptomic atlas allowed us to identify marker genes for each cell population, reconstruct the specification of supporting and steroidogenic cell lineages, to identify the different contributions of coelomic-epithelial cells in male and female gonads and to characterize the expression programs that drive these processes. Our single-cell atlas has been made available via the ReproGenomics Viewer (https://rgv.genouest.org) (Darde *et al*, 2015, 2019).

## Results

### A single-cell expression atlas composed of 128,000 cells from developing fetal gonads

To characterize the cell composition and evolution of human fetal gonads as they differentiate in a sex-specific manner, we collected and dissociated 17 testes and 16 ovaries from 5 to 12 PCW (Fig. 1, panel A). Half of the samples (17 out of 33) originated from young individuals aged from 5 to 7 PCW, so as to be able to decipher early cell differentiation events during and right after sex determination (Hanley *et al*, 2000) (Table S1). The present single-cell atlas is composed of 127,903 cells closely distributed between female (67,427 cells, 53%) and male (60,476 cells, 47%) samples. A principal component analysis of the “pseudobulk” expression matrix showed that the first two components clearly distinguished the samples by gender (35% of the variance of the dataset was explained by PC1) and by developmental stage (26% by PC2) (Fig. 1, panel B) (Cao *et al*, 2019). Samples at 5 PCW, which included emerging gonads together with the underlying mesonephros, consistently segregated from 6-12 PCW gonads that were dissected free of mesonephros. Besides, the close proximity of male and female samples at this earliest stage also suggested that the expression programs that drive sex-specific differentiation of gonadal cells were barely initiated.

**Figure 1.**
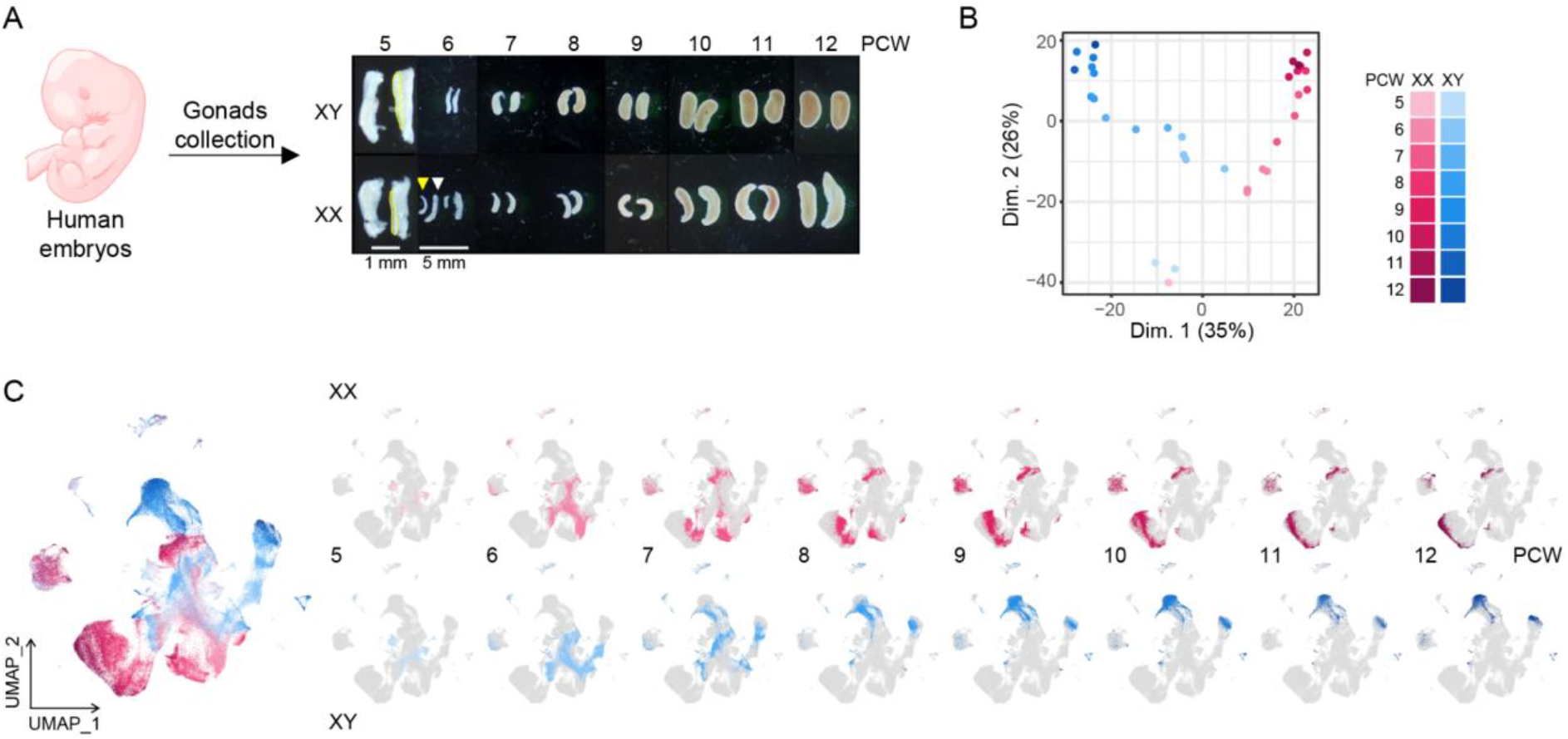
Profiling human fetal gonad development at single-cell resolution. (A) Biological collection for the generation of single-cell RNA-seq data from XX and XY human fetal gonads across eight developmental stages (5 to 12 PCW). At 5 PCW (scale bar = 1 mm), gonads (circled in yellow) were collected together with the underlying mesonephros. From 6 PCW onwards (scale bar = 5 mm), gonads (yellow arrowhead) were dissociated free of mesonephros (white arrowhead). (B) 2D PCA plot of “pseudobulk” RNA-seq profiles showing diverging transcriptomes as XX and XY gonads differentiate from 5 PCW onwards according to the color code. (C) UMAP representation showing all gonadal cells coloured by gender and developmental stage (left). Deconvolution of the global UMAP plot (right) per developmental stage (from 5 to 12 PCW) and gender (XX and XY samples, top and bottom, respectively). In each UMAP plot, all cells from the dataset are in gray while those from a given gender at a specific developmental stage are coloured accordingly.

In accordance with the “pseudobulk” analysis, gonadal cells from individuals of the same developmental stage and gender distributed similarly within the UMAP space (Fig. 1, panel C). Most cells from young developmental stages (5 and 6 PCW) overlapped in the central part of the UMAP, illustrating again the initial transcriptional similarity of XY and XX gonadal cells. Conversely, male and female cells at latter developmental stages occupied outermost and opposite regions of the UMAP, and those from intermediate stages positioned midway between 5 PCW and 12 PCW cells. These distributions imply that cells can be distinguished along developmental stages by expression patterns that change gradually over time and between sex.

### Single-cell transcriptomes distinguish differentiating gonadal cell populations

We next partitioned the 127,903 gonadal cells into 41 clusters (named c1-c41) associated with 20 broad cell types (Fig. 2, panel A; Supplementary results, Section 2) and corresponding to gonadal specific cell lineages as well as common cell types that can be found in most tissues and organs. The expression of known gene markers as well as the percentage of cells belonging to specific gender and/or developmental stage (Fig. 2, panels B and C; Table S2) allowed us to identify : primitive (cluster c32; *HBE1*,+) and definitive (c39; *HBB*+) erythrocytes ; endothelial cells (c35; *CD34*+, *ECSCR*+); immune cells (c26; *PTPRC*+, *TYROBP*+); perivascular cells (c41; *ACTA2*+, *TAGLN*+, *NR2F1*+); primordial (c5, c33; *POU5F1*+, *NANOG*+, *DAZL*+) and female meiotic (c38; *SYCP3*+, *DAZL*+) germ cells; coelomic epithelial cells (c9, c18, c29, c27; *KRT19*+, *UPK3B*+, *LHX9*+); ovarian surface epithelial cells (c10, c22; *LHX2*+, *IRX3*+, *KRT19*+, *UPK3B*+, *LHX9*+); somatic progenitor cells (c3; *IL17B*+, *LHX9*+); interstitial cells (c1, c13, c15, c16, c21, c23, c25; *PDGFRA*+, *TCF21*+, *LHX9*+); fetal/pre-Leydig cells (c27, c34; *INSL3*+, *CYP17A1*+); distinct supporting subpopulations (*WNT6*+) including undifferentiated supporting cells (c2, c24; *DMRT1*+, *TAC1*+, *TSPAN8*+), pre-Sertoli cells (c12; *DMRT1*+, *SRY*+), Sertoli cells (c14, c20, c31; *DMRT1*+, *SOX9*+, *AMH*+), pre-granulosa cells (c6, c7, c8, c11; *FOXL2*+, *IRX3*+); *rete testis/ovarii* cells (c17; *PAX8*+, *TBX1*+). Additionally, several extragonadal cell types (*GATA4*-, *LHX9*-) that originate mostly or exclusively from 5 PCW samples (i.e. gonads dissociated together with the mesonephroi) were captured: non-gonadal coelomic epithelial cells (c28; *UPK3B*+, *KRT19*+, *GATA2*+); mesonephric mesenchymal cells (c19; *PDGFRA*+, *TCF21*+, *NR2F1*+); genital duct cells (c37; *LHX1*+, *PAX8*+, *GATA2*+); neuroendocrine cells (c40; *ASCL1*+, *PHOX2B*+, *GATA2*+); and adrenocortical cells (c36; *NR5A1*+, *CPLX1*+, *NR2F1*+). A small cluster of residual low quality cells (c30) was also identified.

**Figure 2.**
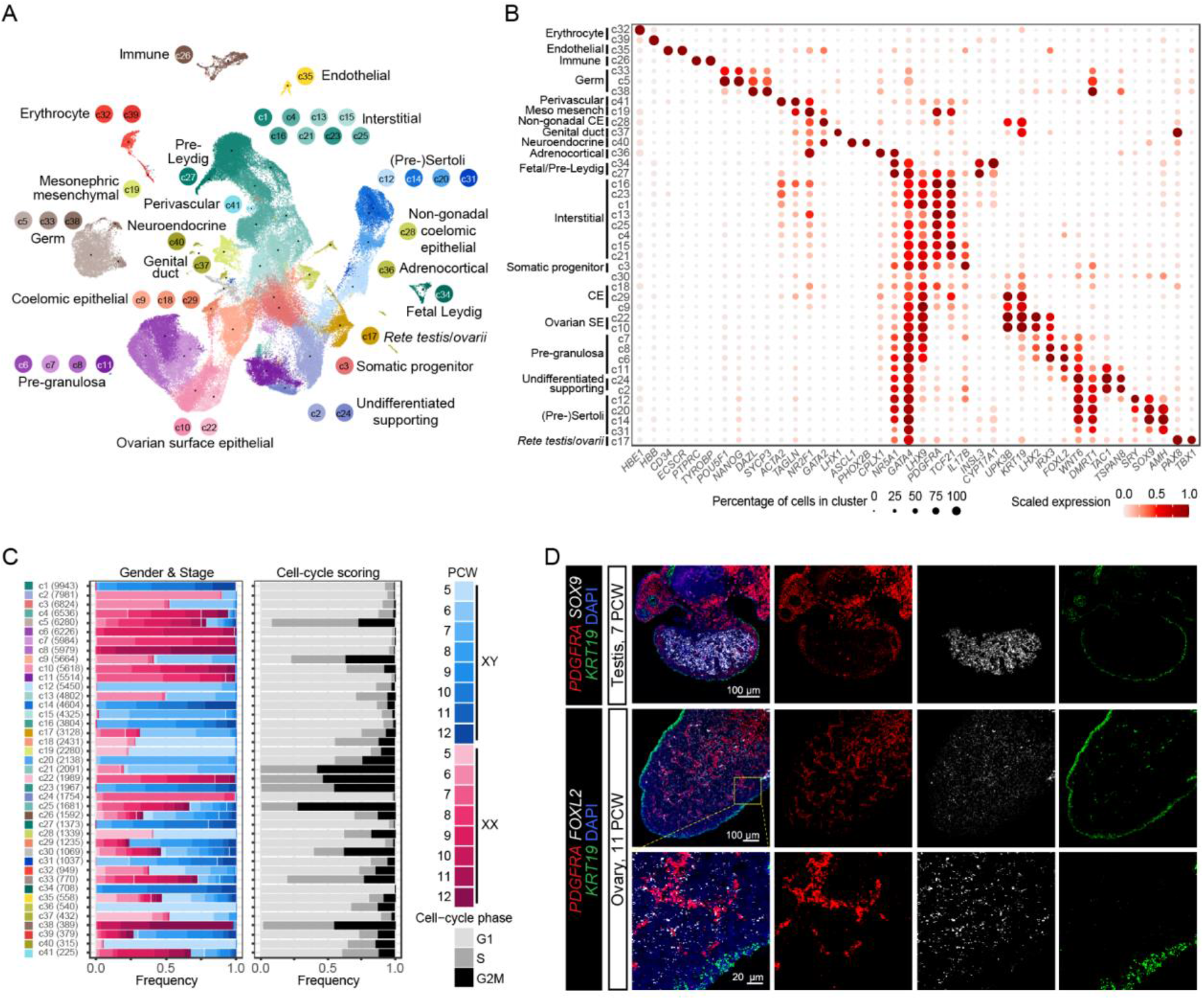
Identification and characterization of the human fetal gonadal cells. (A) UMAP representation of all gonadal cells coloured by cluster (c1-c41) and annotated based on the expression of well-established lineage or cell type markers. (B) Dot plot showing the expression of lineage or cell type markers in each cluster (c1-c41). CE = Coelomic epithelial; SE = Surface epithelial; Meso mesench = mesonephric mesenchymal. (C) Stacked bar plots illustrating the distribution of the gender and developmental stage (left) and of the cell cycle phase assigned to the cells (G1, S or G2/M; right) for each cluster (c1-c41). The number of cells in each cluster is indicated in brackets. (D) RNAscope *in situ* hybridization for gonadal somatic lineage markers, *i.e. KRT19* for coelomic epithelial cells (green), *PDGFRA* for interstitial cells (red), and *SOX9* or *FOXL2* for supporting cells (white), on sections from XY and XX gonads at 7 and 11 PCW, respectively.

Several observations are worth mentioning: first, the exact nature from cluster c3 was initially intriguing as these cells expressed neither supporting nor interstitial markers (Fig. 2, panel B). Considering these cells were almost exclusively observed in early gonads at 5-6 PCW (∼94%), expressed *NR5A1*, *GATA4* and *LHX9* but not *UPK3B* or *KRT19*, together with their central position within the UMAP in-between early interstitial and supporting cells (Fig. 2, panels A-C), we speculated they could represent progenitors of gonadal somatic cell lineages. Second, among male interstitial cells, we identified two clusters related to Leydig cells: cluster c27 and cluster c34, which express higher levels of *INSL3* and of *CYP17A1* respectively, and may represent two functionally distinct subpopulations or, alternatively, two successive steps during their differentiation process (Fig. 2, panel B). As expected, female interstitial cells did not yet express canonical steroidogenic markers. Cluster c4 however was specific to ovaries and is likely to include precursors of theca cells that will arise much later during ovarian development (Fig. 2, panels B and C). Finally, two populations of *PAX8*+ cells could be distinguished: a relatively small cluster of genital duct cells (c37; *LHX1*+, *KRT19*+, *GATA4*-), that were mostly recovered from extragonadal tissue at 5 PCW, and a rather important number of *rete testis* and *rete ovarii* cells (c17; *TBX1*+, *NR5A1*+, *GATA4*+), captured from gonads dissected free of mesonephros from 6 PCW onwards (Fig. 2 panels A-C). 3D imaging on cleared whole-gonads using light sheet fluorescence microscopy indeed confirmed that PAX8 was strongly expressed in Müllerian and Wolffian ducts in the mesonephros as well as in intra-gonadal cells located in the dorsal region of both testes (Fig. 3, panel A) and ovaries (Fig. 3, panel B), nearby the mesonephros. In the testis, the cells expressing PAX8 formed bridges connected to forming testis cords made of Sertoli cells that express SOX9, consistent with the anatomical distribution and organization of the *rete testis* cells (Fig. 3, panel A).

**Figure 3.**
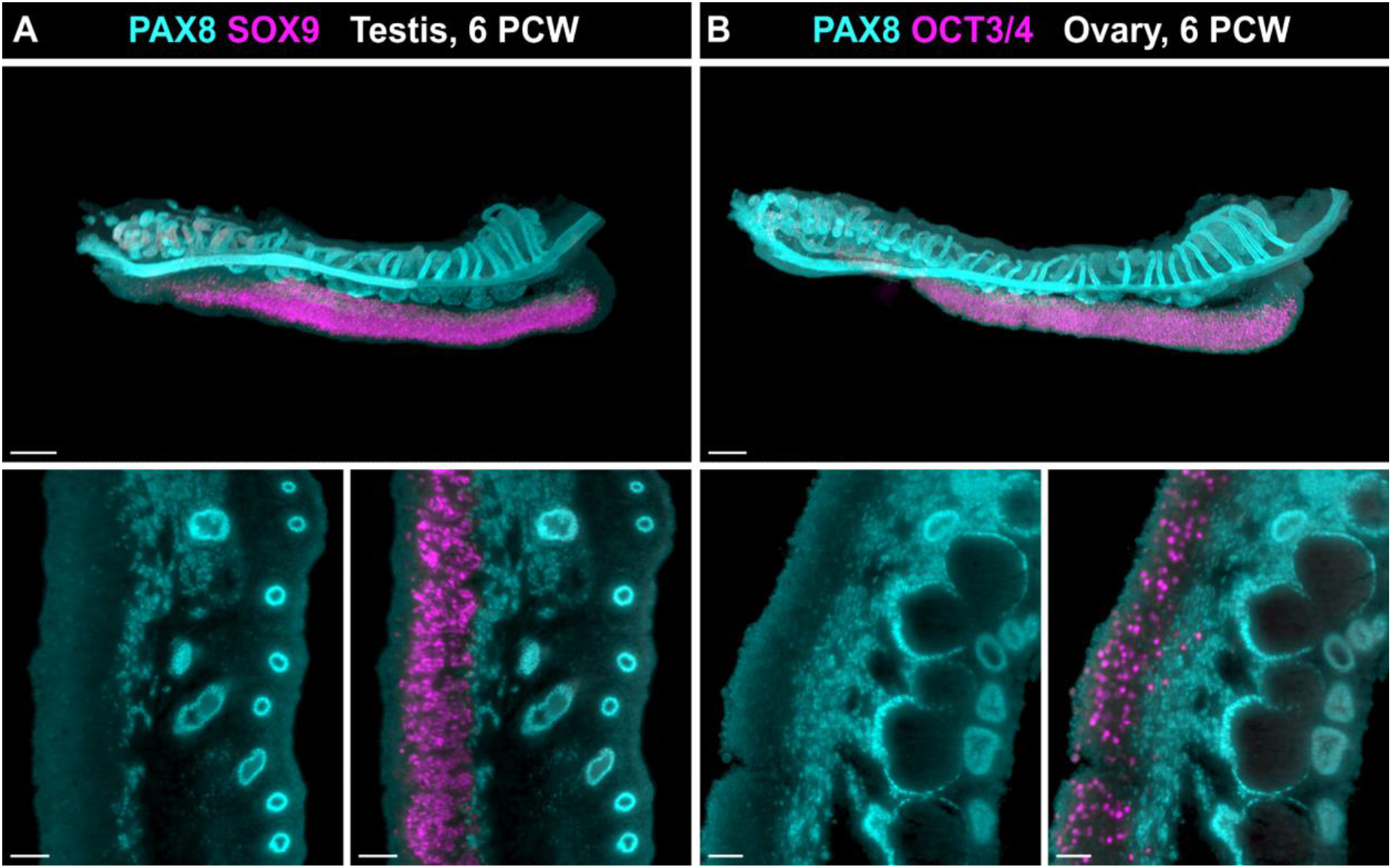
Three-dimensional (3D) imaging of PAX8+ cells in human fetal gonads. (A) 3D visualisation (top panel) of PAX8+ (cyan) and SOX9+ (magenta) cells in an XY fetal gonad at 6 PCW. Two-dimensional (2D) sections (bottom panels) through the gonad and mesonephros highlight intragonadal PAX8+ cells and the continuity with SOX9+ cells within the gonad. (B) 3D visualisation (top panel) of PAX8+ (cyan) and OCT3/4+ (magenta) cells in an XX fetal gonad at 6 PCW. 2D sections (bottom panels) through the XX gonad also reveal a continuity of PAX8+ cells from the mesonephros to the dorsal region of the gonad. Scale bars for top panels = 200µm, for bottom panels = 50µm.

### Disentangling the origin and differentiation of gonadal somatic cells

To tease out the heterogeneity and the differentiation of supporting and steroidogenic cell lineages over developmental stages, we subsetted 111,855 gonadal somatic cells and their putative progenitors which were subsequently partitioned into 42 cell clusters (named s1-s42) (Fig. 4, panels A-D). Among them, 13 clusters consisted of both male and female cells, mostly from 5 to 7 PCW, while the remaining ones were almost exclusively composed of testicular and ovarian cells only (Fig. 4, Panel B). We observed marked changes in terms of cell composition early during male gonad development, especially between 5 and 8 PCW (Fig. 4, Panel C). Such changes on the other hand appeared more progressive in female gonads in agreement with the relative late and slow differentiation process of the ovary as compared to that of the testis.

**Figure 4.**
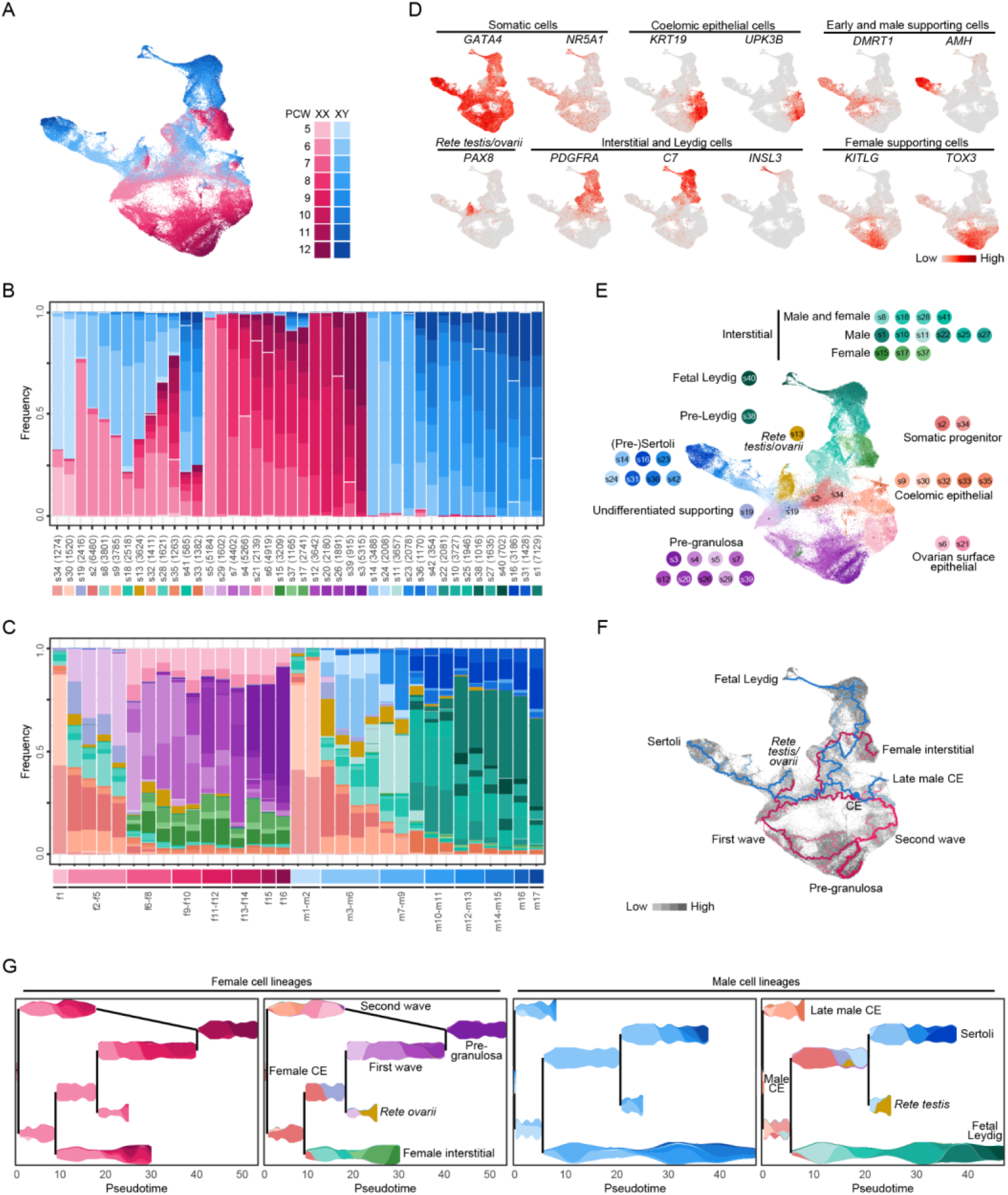
Differentiation of somatic cell lineages during human fetal gonad development. (A) UMAP representation of gonadal somatic cells coloured by gender and developmental stage. (B) Stacked bar plots of the gender and developmental stage in each cluster (s1-s21), with the number of cells for each cluster indicated in brackets. (C) Stacked bar plots of the cell cluster in each of the 33 samples. (D) UMAP visualization of the scaled expression of gene markers in gonadal somatic cells. (E) UMAP representation of gonadal somatic cells coloured by cluster (s1-s21) and annotated by cell population based on the expression of established lineage or cell type markers. (F) Gender-specific trajectories of gonadal somatic lineages. Starting points of each trajectory are indicated as circles located in cluster s30. Pink and blue lines correspond to female and male trajectories, respectively, while cells are coloured from light to dark gray according to increasing pseudotime. (G) Visualization of female and male gonadal somatic cell lineages (left and right, respectively). Cell density along pseudotime (x-axis) within each branch of the lineages is presented using violin plots that are coloured either by gender and developmental stage (left) or by cell cluster (right), with cell fate decisions visualized as branch points. CE = Coelomic epithelial.

We next investigated the sex-specific developmental trajectories that fetal somatic subpopulations follow as human gonads differentiate. As a prerequisite to interpreting such trajectories we first used known marker genes to annotate the 42 somatic cell clusters (Fig. 4, panels D and E, Table S3). We then performed pseudotemporal ordering of XX and XY somatic cells separately using Monocle3. Since gonads originate from the coelomic epithelium at the surface of the mesonephros, we used the medoid of coelomic epithelial cells (i.e. cluster s30) at 5 PCW from each sex as starting points, yielding eight major trajectories (Fig. 4, panels F and G). Strikingly, the identified trajectories were highly consistent with developmental stages, both male and female cells being ordered with increasing pseudotime from 5 to 12 PCW. Initial trajectories for male and female cells were highly overlapping (Fig. 4, panel F): disregarding their sex, early coelomic epithelial cells (cluster s30) first evolved through cluster s34 and up to cluster s2, prior to bifurcating towards either the interstitial lineage or the supporting lineage. The specification of these two lineages in both males and females from a unique cell subpopulation suggests that cells from cluster s2 are the multipotent somatic progenitor cells derived from the coelomic epithelium. Of note is that most cells from cluster s2 (87,6%) belonged to cluster c3 in the global analysis (Fig. 2, panel A), which we already hypothesized to correspond to gonadal progenitors. The initial commitment of XX and XY somatic progenitors to the interstitial fate involved similar trajectories through cluster s8 and s18, prior to acquiring gender-specific paths. These included clusters s10, s22, s1 and s38 up to the fetal Leydig cell (s40) for testicular cells, and clusters s37 and s17 up to s15 for ovarian cells (Fig. 4, panel F). Similarly, initial trajectories that reflect the specification of male and female supporting lineages overlapped up to cluster s19 but quickly diverged towards two sexually-dimorphic early supporting subpopulations: cluster s24 in testes and cluster s5 in ovaries, which differentiate further into Sertoli cells up to clusters s31 and s16 and pre-granulosa cells up to clusters s39 and s3, respectively (Fig. 4, panel F). Overall, these trajectories recapitulate the widely accepted model in which coelomic epithelium-derived progenitors give rise to both supporting and interstitial lineages that rapidly commit to one or the other of their sex-specific fates (Fig. 4, panel G).

In addition to this canonical model, our analysis also identified branching points along the supporting lineage of both male (cluster s24) and female (cluster s5) gonads. The two resulting secondary trajectories lead to the *rete testis* and *rete ovarii* cells (cluster s13; Fig. 4, panels D, F and G), indicative of a supporting origin of these cells as recently suggested in the mouse (Kulibin & Malolina, 2020; Mayère *et al*, 2021b). Furthermore for XX gonads we observed a second trajectory that originated from ovarian surface epithelial cells (clusters s21 and s6) and led to the formation of pre-granulosa cells (clusters s39 and s3; Fig. 4, panel D). Importantly these two trajectories converged at a pseudotime composed of cells from cluster s20 at 9 PCW, therefore making a loop towards a common pathway for the differentiation of most advanced pre-granulosa cells at 12 PCW (Fig. 4, panels F and G). It is thus tempting to speculate that this alternative pathway actually corresponds to the specification of cortical granulosa cells directly from the ovarian surface epithelium. In contrast, coelomic epithelial cells in developing testes only evolved towards a slightly different state (s33) at the later stages of this study.

### Cell trajectory inference deciphers pseudotemporal gene expression analysis during differentiation of supporting and interstitial cell lineages

To highlight potential key regulators driving somatic cell specification and differentiation in human fetal gonads, we focused on the 4,387 differentially-expressed genes (DEGs) with dynamic expression over the eight major cell trajectories. Among these, 187 genes are known to be important or involved in sex determination, gonad development and/or steroidogenesis, while 113 other ones encode for transcription factors that could be required for or involved in the establishment of gonadal somatic cell lineages (Table S4). We further classified the DEGs into 24 co-expression modules (termed m1 to m24) according to their overall expression patterns in distinct somatic cell lineages (Fig. 5, panel A and Table S4). The modules revealed waves of expression with induction along each female or male lineage, starting with the coelomic epithelial cells at early pseudotime and developmental stages, and followed by the female and male supporting cells, the *rete ovarii*/*testis* and the interstitial cells (Fig. 5, panel A). Four modules corresponded to genes that were already expressed in early coelomic epithelial cells (m1-m3 and m17). These genes were subsequently downregulated in interstitial cell lineages (m1; *GATA4*, *NR0B1*, *RSPO1*, *WT1*), in supporting cell lineages (m17; *SFRP1*, *LHX9*, *WNT5A*), in all somatic cell lineages (m2; *HMGA1*, *WNT2B*) or were further upregulated in late coelomic epithelial cells (m3; *BNC1*, *KRT19*, *UPK3B*). A single module consisted of genes showing peak expression in gonadal somatic progenitor cells (m4; *KDR*, *CCL21*, *GEM*, *IL17B*), prior to the differentiation of all supporting and interstitial cell lineages with the exception of the second wave of pre-granulosa cells that directly arise from the ovarian surface epithelium (Fig. 5, panel B). Regarding genes with preferential expression within supporting cell lineage, eleven modules could distinguish those induced in both female and male supporting cells (m5-m8), specifically in pre-granulosa cells (m9-12) or in (pre-)Sertoli cells (m13-m15). More specifically, some genes showed a prolonged (m5; *GADD45G*, *WNT6*) or transient (m6; *DMRT1*, *GFRA1*, *REG3G*, *TAC1*, *TSPAN8*) expression in all supporting cell lineages except during the second wave of pre-granulosa cells, while others were also activated during the second wave of pre-granulosa cells, again in a prolonged (m7; *TAF4*, *NRIP1*) or transient (m8; *CYP26B1*, *RUNX1*) manner. Concerning genes expressed more specifically in pre-granulosa cells, four modules were identified and included genes induced early (m9; *ESR1*, *KITLG*, *IRX3*, *TOX3*) or late (m10; *FOXL2*, *YBX3*) during the two differentiation waves, only during the second wave (m11; *LHX2*, *POU6F2*) or only during the first wave (m12; *CYP19A1*, *GHRL*). On the other hand for male supporting cells, three modules of genes that showed early (m13; *SRY*), intermediate (m14; *DHH*, *FATE1*, *INHBB*, *SOX10*, *SOX9*) or late (m15; *AMH*, *CITED1*, *NKX3-1*) induction during the differentiation of Sertoli cells were identified. Module m16 corresponded to genes with peak expression in *rete ovarii* and *rete testis* cells (m16; *FBXO32*, *PAX8*, *TBX1*). In addition to module m17 (see above), several modules were associated with distinct expression patterns within the interstitial cell lineages (m18-m22). These included genes expressed throughout the entire interstitial lineage of both male and female gonads (m18; *ARX*, *HIC1*, *PDGFRA*, *TCF21*) or more sequentially and transiently, with early (m19; *RXFP2*), intermediate (m20; *CYP1B1*, *NUPR1*) or late and male-biased (m21; *INHBA*, *MSC*, *OSR2*, *PTCH1*, *PTCH2*) induction. Finally, one module included genes with peak expression at the end of the male interstitial cell lineage, in Leydig cells (m22; *CYP17A1*, *INSL3*, *FOXO4*, *LHCGR*, *NR5A1*). Noteworthy is that several of these genes involved in steroidogenesis and expected to be specifically expressed in Leydig cells were also found in early supporting and/or (pre-)Sertoli cells (e.g. *CYB5A*, *CYP11A1*, *CYP17A1*, *HSD3B2*, *STAR*), albeit at lower levels. This suggests a potential functional interplay between Sertoli and Leydig cells for the synthesis of androgens during fetal life, as already proposed in the mouse (Shima *et al*, 2013). These expression profiles are also consistent with the dynamics of *NR5A1* that was found preferentially expressed in Sertoli cells at 6-7 PCW, prior to being enriched in Leydig cells from 8 PCW onwards (Fig. S1). Finally, two modules included genes from the Y chromosome (m23), that were expressed only in male lineages, and genes involved in cell cycle (m24), that were expressed in coelomic epithelial cells and transiently within the interstitial cell lineages.

**Figure 5.**
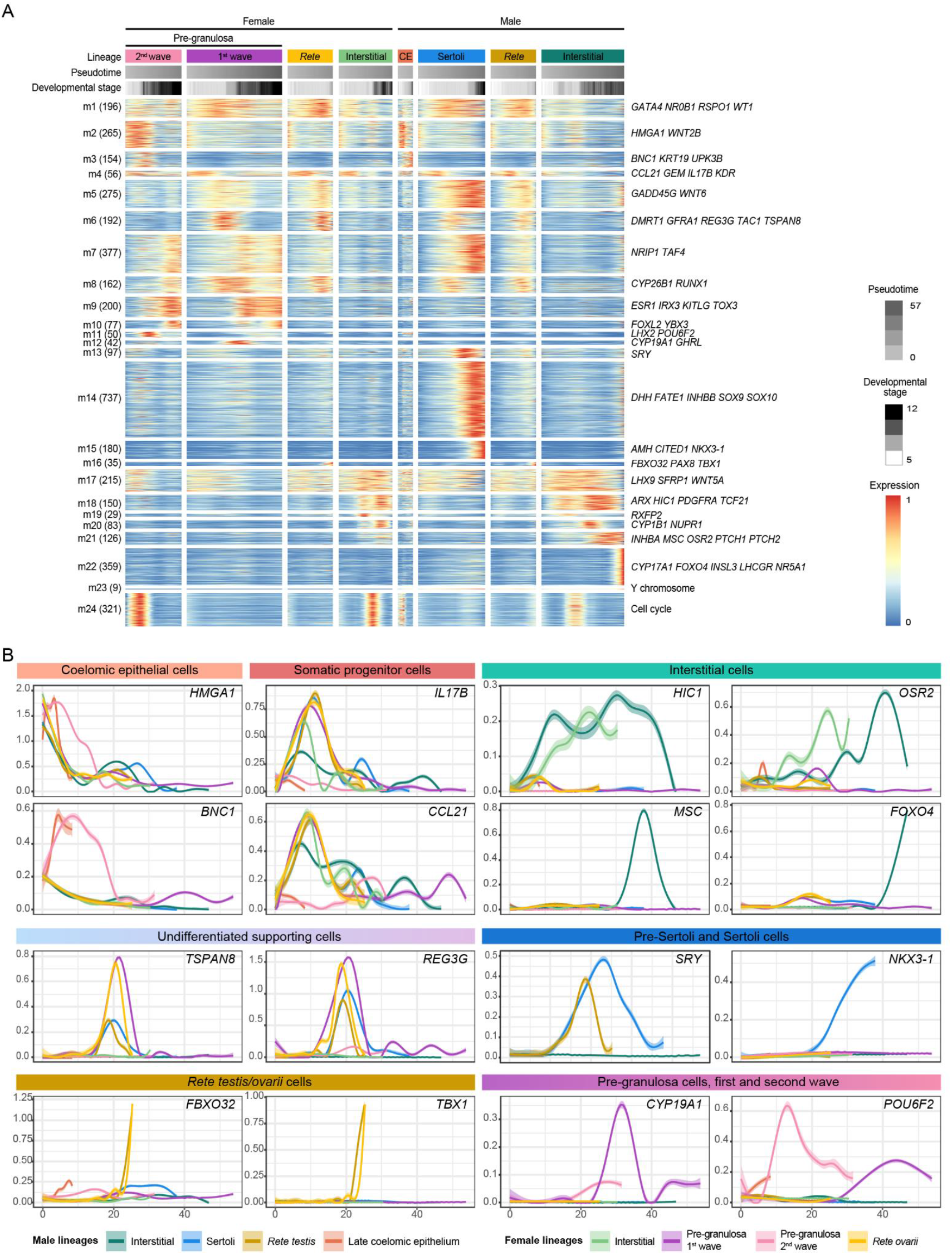
Expression dynamics during gonadal somatic cell differentiation. (A) Heatmap illustrating modules (m1-m24) of coexpressed genes along the pseudotime in female (left) and in male (right) lineages. For each lineage, horizontal bars illustrate the pseudotime and the developmental stage according to the color codes. The number of differentially-expressed genes is indicated in brackets at the right of the module identifier and some associated markers are indicated at the right. A color scale for scaled expression data is given. (B) Gene expression profiles smoothed over pseudotime and coloured by cell lineage. CE = Coelomic epithelial.

Next, we investigated further the spatial distribution of supporting cells thanks to newly identified markers: *WNT6* (module m5) for all supporting cells, *TAC1* (m6) for early supporting cells, and *LHX2* (m11) for pre-granulosa cells derived from the coelomic epithelium (Fig. 6, panel B). We first validated *WNT6* as a supporting cell marker for both male and female gonads, as evidenced by its coexpression with *SOX9* and *FOXL2*, respectively, on top of *WT1* (Fig. S2). Consistent with its early expression in undifferentiated supporting cells, we detected *WNT6* in a similar pattern to that of *SRY* prior to cord formation in the testis at 5 PCW and before the onset of *FOXL2* expression in the ovary at 6 PCW (Fig. 6). On the other hand *TAC1* was detected mostly in outer male supporting cells at 5 PCW, in a complementary manner to *SOX9* that was found to be preferentially expressed in inner supporting cells. While *TAC1* expression sharply decreased in testes from 6 PCW onwards (Fig. S3), it was detected at high levels in inner supporting cells of ovaries, at least up to 11 PCW (Fig. 6, Fig. S4 and data not shown). Strikingly, *FOXL2* also showed a gradient expression with decreasing levels from center to periphery, albeit with a broader expression pattern than that of *TAC1* (Fig. 6 and Fig. S4). Conversely, *LHX2* was strongly detected in female coelomic epithelial cells and in outer pre-granulosa cells, with decreasing expression levels from periphery to center of ovaries (Fig. 6, Fig. S4 and Fig. S5).

**Figure 6.**
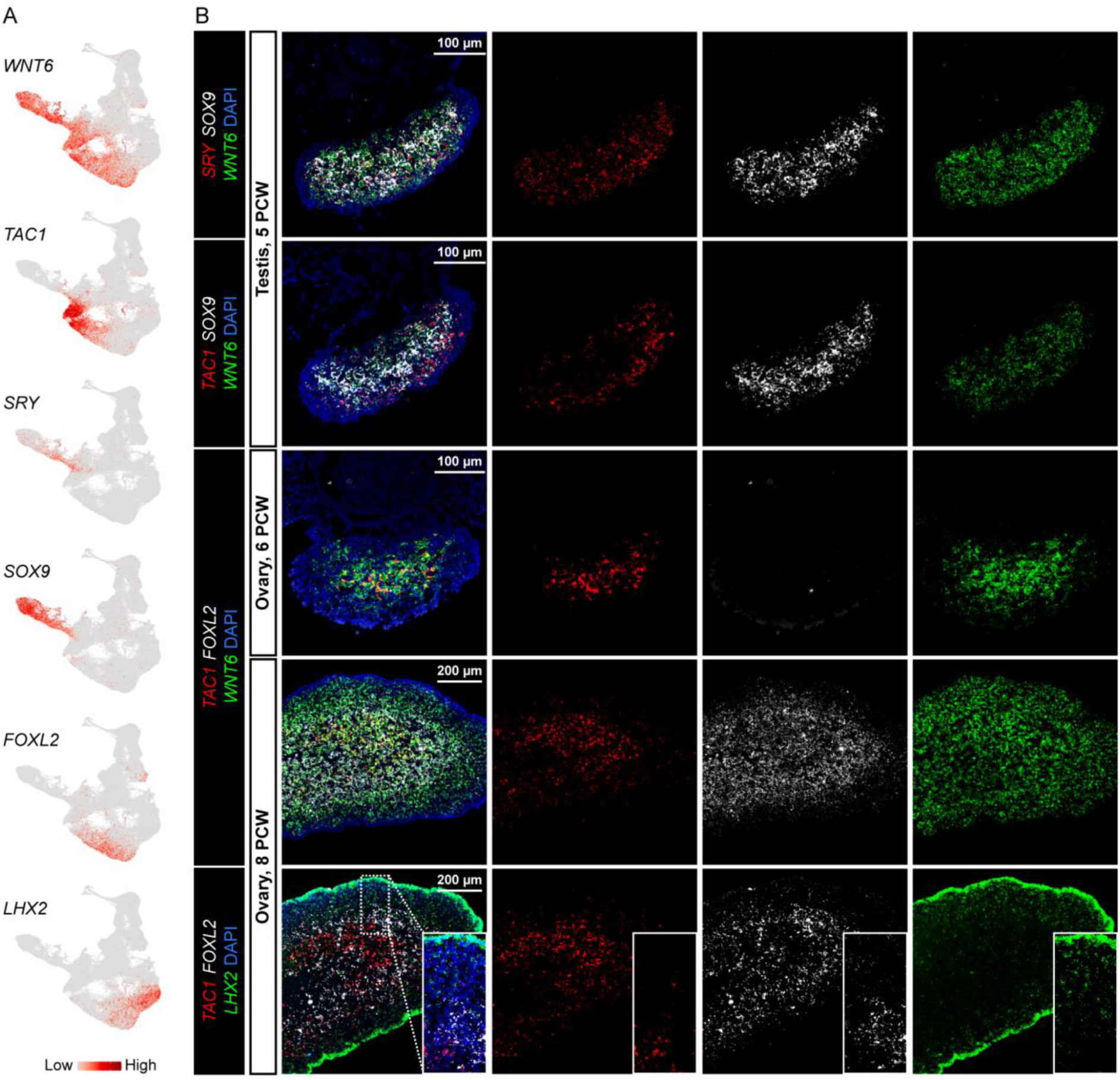
Spatial localization of supporting cells in human fetal gonads. **A.** UMAP visualization of the expression of *WNT6* (Supporting cells), *TAC1* (Early supporting cells), *SRY* (Pre-Sertoli cells), *SOX9* (Sertoli cells), *FOXL2* (Pre-granulosa cells) and *LHX2* (Ovarian surface epithelial cells and pre-granulosa cells). **B.** RNAScope *in situ* hybridizations for *SRY* or *TAC1* (red), *SOX9* or *FOXL2* (white), *WNT6* or *LHX2* (green) on sections from a XY gonad at 5 PCW and XX gonads at 6 and 8 PCW; the short side of the inset (bottom panel) is 100 µm.

Altogether, the combined trajectory and differential expression analyses finely recapitulated the transcriptional programs that distinguish differentiating somatic cell lineages across time, sex and space in human fetal gonads (Fig. 7).

**Figure 7.**
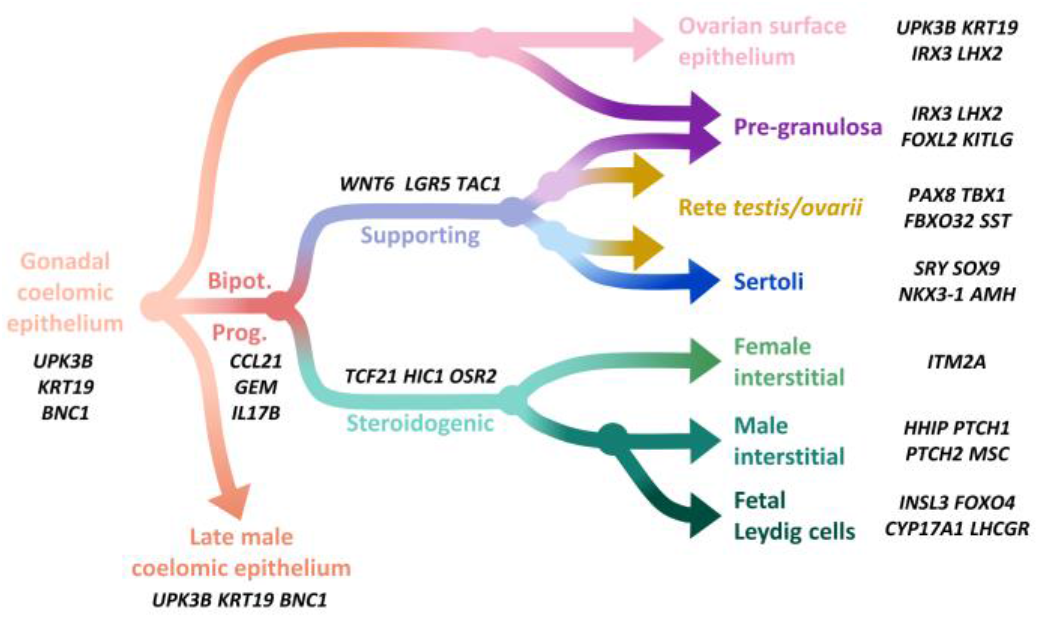
Specification and differentiation of somatic cell lineages in first trimester human fetal gonads. The cell lineages derived from the gonadal coelomic epithelium are presented together with associated gene markers.

## Discussion

Our current understanding of human sex differentiation is hampered by an incomplete characterization of all gonadal cell types and their expression programs, especially at early stages. Indeed, most of the knowledge on human gonad development comes from samples mostly older than 7 PCW. Furthermore, several bulk transcriptomic analyses were conducted to gain insights into the gene networks underlying human gonad differentiation (Del Valle *et al*, 2017; Lecluze *et al*, 2020). Although these approaches highlighted developmentally-regulated and sexually dimorphic genes, they failed to unravel the cellular heterogeneity of human fetal testes and ovaries. Yet, the identification and characterization of gonadal somatic cells at each step of their differentiation process, including the progenitor cells, is compulsory to understand how distinct cell lineages are specified and progress during sex determination. Here we leveraged the power of single-cell RNA-sequencing to study cell fate determination during gonad differentiation in the first trimester of human development, from 5 to 12 PCW.

### An atlas of gonadal sex determination at the single-cell level in humans

For this study we captured and sequenced individual cells from 33 male and female gonads that covered a developmental window of seven weeks from sex determination onwards. While a strong focus was made to densely and finely characterize cell composition of undifferentiated gonads, our experimental design also allowed us to recover more advanced cell differentiation states during gonad development. Altogether our dataset contains most, if not all, expected cell types of the gonad, including red cells, immune cells, endothelial cells and perivascular cells, on top of germ cells, supporting cells and interstitial cells at distinct differentiation stages, as well as their presumptive multipotent progenitors. We characterized gene expression in each of these populations and identified new markers, including human-specific ones, and made this atlas available to the community through an online viewer. Several studies also investigated human gonad development at the single-cell level. These were however focused on gonads from one sex or the other, included only few samples at early stages and/or no biological replicates (Chitiashvili *et al*, 2020; Wang *et al*, 2022; Guo *et al*, 2021). More recently, one exhaustive study reported the single-cell transcriptomic analysis of 38 human gonads during the first and second trimester of development (Garcia-Alonso *et al*, 2022). Again, very few samples younger than 8 PCW were included, and most samples included extragonadal tissue, making the use of spatial transcriptomics needed to eventually reattribute all captured cells to their tissue of origin. Overall, the dataset described here represents to our knowledge the most comprehensive single-cell transcriptomics resource of early human gonad development.

### Origin and specification of gonadal somatic cell lineages

Beyond building an atlas of gonad development, our main objective was to explore the differentiation pathways of somatic cells in early human fetal gonads. Using trajectory analyses, we showed that the specification of human gonadal somatic cell lineages follows the dogma of sex determination as established in the mouse. Indeed, gonadal progenitors, derived from the coelomic epithelium, can give rise to both the interstitial and supporting cell lineages that will subsequently commit to sex-specific fates. Nevertheless, we found that the expression programs that underlie the specification of these lineages are not necessarily conserved in the mouse. This is notably striking for early supporting cells that specifically express strong levels of *e.g. TAC1* or *TSPAN8*, which are not expressed in the mouse gonads (Niu & Spradling, 2020; Garcia-Alonso *et al*, 2022). Importantly, we also characterized the phenotype of human gonadal progenitors, whose cell population is almost exclusively present in male and female gonads at 5 and 6 PCW. These progenitors lose the coelomic epithelial cell markers *UPK3B* and *KRT19*, they maintain the expression of the gonadal somatic cell markers *NR5A1* and *GATA4*, and they co-express markers characteristics of the interstitial and the supporting lineages such as *TCF21*, *PDGFRA*, *LHX9*, *DLK1*, *NPY* and *SPRR2F*. Additionally, they display a preferential expression for *GEM*, *IL17B*, *CCL21* and *KDR*. Together with coelomic epithelial cells (44-58% of all gonadal somatic cells), progenitor cells constitute the most abundant population (37-44%) within 5 PCW gonads. At 6 PCW, their proportion drops to 17% and 7% in female and male gonads, respectively. At this stage, early supporting cells represent 21 to 54% of somatic cells, and already include *SOX9*+ Sertoli cells in the testis, demonstrating that sex differentiation is initiated earlier than previously described (Yang *et al*, 2018; Wang *et al*, 2022). These results also suggest that sex differentiation in humans, especially when compared to the mouse, is a relatively slow and asynchronous process, since supporting and interstitial cells at distinct differentiation stages could be found in the same gonad together with progenitor cells. Such asynchrony also clearly highlights the higher relevance of single-cell approaches over bulk experiments to study developmental processes.

### Evolution of the gonadal sex differentiation dogma

The main strength of single-cell genomics resides in its ability to capture and characterize cells without prior knowledge, enabling the identification of rare or previously unknown cell types and states. While being known at anatomical and histological levels, the *rete testis* and *rete ovarii* cells for instance have remained poorly explored and have been somehow excluded from the sex differentiation process for decades. A few years ago, these cells were shown to harbor supporting-like features and to express PAX8 in the mouse (Malolina & Kulibin, 2019; Kulibin & Malolina, 2020). More recently, scRNA-seq analysis of fetal mouse gonads identified the *PAX8*-expressing cell population as being the first lineage to emerge from the coelomic epithelium, before the specification of supporting cells, while lineage tracing experiments further suggested these cells contribute to the pool of male and female supporting cells (Mayère *et al*, 2021b). We and others (Garcia-Alonso *et al*, 2022; Taelman *et al*, 2022) also characterized this not so new cell population in the human fetal gonad, with PAX8 representing an evolutionary-conserved marker for this *rete* lineage. Unlike in the mouse, our data suggest that human *rete* cells differentiate from early supporting cells rather than directly from the coelomic epithelial cells. Whether PAX8-expressing cells could also have the capacity to give rise to supporting cells in humans remains to investigate. Our analyses also highlight for the first time two distinct populations of Leydig cells that differentially express *INSL3* and steroidogenic genes. The coexistence of these two populations from around 8 PCW up to at least 12 PCW raises several questions regarding their origins, differentiation mechanisms and fates. It could be important for instance to investigate whether these populations actually arise from two distinct progenitor lineages as described for mouse fetal Leydig cells (DeFalco *et al*, 2011). We believe that this is not the case, as these two populations express *GATA4* and *NR51A1*, but rather that they represent two sequential states during Leydig cell differentiation, as suggested by trajectory analyses. It will also be interesting to evaluate whether signaling mechanisms known to regulate the specification of mouse Leydig cells such as the PDGFRA (Brennan *et al*, 2003) or the DHH (Yao *et al*, 2002) pathways may also trigger the differentiation of one and/or the other of these two Leydig cell populations in humans. One might also hypothesize that one of these populations could represent progenitors of adult Leydig cells, whose origin is still heavily debated (for review, see (Shima, 2019)), while the other population would represent fetal Leydig cells intended to involute later during development. Further investigations are therefore needed to answer these questions, within the limits of available experimental procedures for human tissues.

### Distinct spatio-temporal pathways for the differentiation of supporting cells

The inclusion of an important number of early gonads in our study allowed us to finely characterize the initial steps of supporting cell differentiation. We notably identified strong, human-specific markers for early supporting cells, including *TAC1*, *TSPAN8* or *REG3G*. All of them are expressed relatively transiently, right after the somatic progenitor step and before the onset of the canonical markers *SOX9* and *FOXL2*. A recent study proposed *TAC1+* cells to be exclusively found in ovaries and to represent pre-granulosa cell progenitors (Wang *et al*, 2022).In our dataset we clearly detected *TAC1* in all male gonads at 6 PCW. Moreover, we found *TAC1* to be strongly expressed in *SRY*+/*SOX9*-pre-Sertoli cells at the periphery of the testis at 5 PCW, while it was expressed at lower levels in *SRY*+/*SOX9*+ Sertoli cells which were located deeper. This clearly indicates that *TAC1*+ cells represent an early step during supporting cell specification rather than an “ovarian-specific” progenitor population. This spatial distribution further suggests that the differentiation of male supporting cells proceeds from the center to the periphery of the testis, with first-specified pre-Sertoli cells being surrounded by newer ones. Such gene expression gradients were also reported for *SOX9*, *DMRT1* and *GSTA2* in chicken fetal testes (Estermann *et al*, 2020b). In ovaries, we detected *TAC1* in *WNT6*+/*FOXL2*-pre-granulosa cells at 6 PCW as well as in *WNT6*+/*FOXL2*+ pre-granulosa cells at least up to 11 PCW. Nevertheless, unlike in the testis, *TAC1* was restricted to the innermost, medullary pre-granulosa cells and was absent in the outermost, cortical pre-granulosa cells, which expressed *LHX2* instead. The specific locations of these distinct female supporting cell populations is consistent with our lineage analysis that showed two waves for the differentiation of pre-granulosa cells: a first one that initiates from the coelomic epithelium-derived somatic progenitors, which also gives rise to pre-Sertoli cells in male gonads, and a second one that derives directly from late female coelomic epithelial cells. This developmental scheme appears similar to the mouse model where *Foxl2*+ medullary pre-granulosa cells were described to emerge from *Wnt6*+ bipotential progenitors while *Lgr5*+ cortical pre-granulosa cells were shown to derive from *Krt19*+ epithelial progenitors (Niu & Spradling, 2020; Rastetter *et al*, 2014). However, in contrast to the mouse, we found that *LGR5* was expressed in early supporting cells derived from somatic progenitors in both male and female gonads, and not in cortical pre-granulosa cells derived from coelomic epithelial cells (Fig. S4).

## Conclusion

Our single-cell transcriptomic analysis provides a comprehensive understanding of how male and female gonadal cells differentiate during sex determination in humans. In addition to the characterization of new cell subpopulations within the gonads, we uncovered the expression programs that underlie the commitment of distinct somatic cell lineages from their common progenitor cells. Notably, our single-cell atlas enriched in young individuals aged from 5 to 7 PCW allowed us to decipher early gonadal cell specification events and to identify associated markers, including for progenitor cells. Furthermore, and although localization is necessarily lost in single-cell experiments, we resolved the molecular identity of supporting cells that arise from distinct origins and distribute in different regions of the human fetal ovary. Awaiting for the improved resolution of spatially resolved transcriptomics technologies, the new markers we identified here also represent promising tools to accurately investigate the morphological relationship between gonadal cells using dedicated RNA- or protein-based technologies. Additionally, our study paves the way towards a systematic cross-species comparison of gonadal sex differentiation, which will be instrumental to highlight those molecular factors and signaling pathways that would deserve specific attention to understand gonad development and related disorders in humans.

## Material and methods

### Ethic statement

First trimester human embryos and fetuses (5-12 PCW) were obtained from legally induced terminations of pregnancy performed in Rennes University Hospital. No termination of pregnancy was due to fetal abnormality. Tissues were collected following women’s written consent, in accordance with the legal procedure agreed by the National agency for biomedical research (authorization #PFS09-011; Agence de la Biomédecine) and the approval of the Local ethics committee of Rennes Hospital (advice # 11-48).

### Sample collection

The terminations of pregnancy were induced using a standard combined Mifegyne® (mifepristone) and Cytotec® (misoprostol) protocol, followed by aspiration. A subset of women received either 400 or 800 mg of ibuprofen for preventive analgesia or none. Gestational age was determined by ultrasound, and further confirmed by measurement of foot length (Evtouchenko *et al*, 1996; O’Shaughnessy *et al*, 2019). The samples were recovered from the aspiration products using a binocular microscope (Olympus SZX7, Lille, France) and were immediately placed in ice-cold phosphate-buffered saline (PBS). For 5 PCW samples, gonads and the underlying mesonephros were collected together while from 6 PCW onwards gonads were dissected free of mesonephros. The sex of individuals was determined by morphological evaluation of the gonads, except for embryos younger than 7 PCW, for which a qPCR was performed on genomic DNA using primers specific for SRY (Forward: ACAGTAAAGGCAACGTCCAG; Reverse: ATCTGCGGGAAGCAAACTGC) (Friel *et al*, 2002) and for a genomic upstream of the GAPDH gene (Forward: CCACAGTCCAGTCCTGGGAACC; Reverse: GAGCTACGTGCGCCCGTAAAA) (Du *et al*, 2015).

### Tissue dissociation

Single-cell RNA-sequencing was performed on human fetal gonads from 33 individuals (16 females and 17 males) ranging from 5 to 12 PCW. The 5 PCW samples included n=1 female and n=2 males. Then, for each sex, samples included n=4 at 6 PCW, n=3 at 7 PCW, n=2 at 8 PCW, n=2 at 9 PCW, n=2 at 10 PCW and n=2 at 11-12 PCW (Table S1). Single cell suspensions were obtained by a standard enzymatic and mechanical digestion procedure. Briefly, gonads were cut into small pieces and digested in PBS containing 0.25% Trypsin-0.02% EDTA (#T4049, Sigma-Aldrich) and 0.05 mg/ml DNase (#DN25, Sigma-Aldrich) for 10 min at 37 °C, followed by gentle up and down pipetting. Trypsin digestion was stopped by adding 10% fetal bovine serum in M199 medium and centrifugation at 350 g for 5 min at 37°C. Dissociated cells were resuspended and washed twice in PBS containing 0.04% weight/volume BSA. The final cell suspensions were filtered on 30 µm strainers (Miltenyi Biotec) and counted on a Malassez hemocytometer after Trypan blue staining.

### Single-cell capture, library preparation and sequencing

All steps for the construction of single-cell libraries (Chromium™ Single Cell 3’ Library & Gel Bead Kit v2) were carried out according to the manufacturer’s instructions (10x Genomics). In brief, cell concentration was systematically adjusted to 400 cells/µL in PBS-BSA prior to loading on the Chromium™ Single Cell A Chip. For each sample, ∼7,000 cells were loaded on a single reaction well in order to target ∼4,000 recovered cells. Following construction, the 33 single-cell libraries were quality-controlled using a bioanalyzer (Agilent Technologies), pooled and sequenced on a single lane of an Illumina HiSeq 4000 instrument using a 2×100 paired-end sequencing protocol (Illumina, San Diego, CA) to validate quality and to adjust library pooling according to actual cell capture rates. Subsequently, the 33 pooled libraries were sequenced on 33 lanes of an Illumina HiSeq 4000 instrument to achieve an average sequencing depth of at least 350 million paired-reads per sample.

### scRNA-seq data pre-processing, quality controls and normalization

Demultiplexed raw sequencing reads were processed, mapped to the GRCh38 human reference genome (official Cell Ranger reference, v3.0.0) and quantified based on unique molecular identifiers (UMIs) with the Cell Ranger pipeline (v7.0.0, 10x Genomics). The resulting raw count matrices from the 33 individual samples were then merged without normalization (Cell Ranger v7.0.0). The cell-calling algorithm from Cell Ranger, which relies on the EmptyDrops method (Lun *et al*, 2019), was also used to establish a first set of potential valid cells.

Subsequently, cells with fewer than 500 detected genes; fewer than 2000 counts or higher than 100,000 counts; more than 30% of mitochondrial genes; or detected as doublets by the DoubletFinder V3 R package were removed from the aggregated matrix. Cell cycle phases were assigned using the Seurat (v4.0.1) R package. Finally, the data were normalized and scaled using the SCTransform function with cell-cycle score and mitochondrial-encoded gene transcript levels as covariates.

“Pseudobulk” samples (log2-transformed and normalized count matrix from the aggregated counts from all the analyzed cells of each sample) were also projected on a 2D PCA-based space using the FactoMineR package to graphically evaluate their distribution (Fig. 1, panel B).

Regarding the focused analysis of gonadal somatic cells, we first selected relevant clusters identified from the primary analysis and further reanalyzed the 111,855 subsetted cells with a similar strategy.

### Reduction of dimensionality and clustering

Variable genes were detected using the SCTransform function from Seurat (v4.0.1) and used to perform a PCA reduction of dimensionality. The first 50 principal components were then used in the Louvain graph-based clustering using the FindNeighbors function followed by the FindClusters functions with a resolution of 1.2 and 1.6 for the initial analysis and for the somatic cell analysis, respectively. Finally, we used the uniform manifold approximation and projection (UMAP) method implemented in the Seurat package to project single cells in a reduced 2D space.

### Cluster annotation and differential gene expression analysis

Cell clusters and subpopulations were annotated using known marker genes as well as computed differentially expressed genes using the Wilcox rank sum test implemented in Seurat package (v4.0.1) (one-tailed Wilcoxon rank sum test, p-values adjusted for multiple testing using the Bonferroni correction).

### Pseudotemporal cell trajectory inference

To order gonadal somatic cells in pseudotime and infer male or female trajectories over developmental stages, we used the learnGraph and orderCells functions using default parameters (except for learnGraph: close_loop = loop) implemented in the Monocle3 R package (version 1.0.0) (Cao *et al*, 2019). To specify the starts of the male and female lineage trajectories and to smooth expression values over pseudotime, we first selected XY and XX coelomic epithelial cells (cluster s30) at the earliest developmental stage (5 PCW), respectively. The position of the root node of each trajectory was then determined by extracting the closest neighboring nodes of the selected cells.

Differentially expressed genes over each of the eight major pseudotemporal cell trajectories were determined with the tradeSeq package with a number of knots (nknots) = 6 (Van den Berge *et al*, 2020). Briefly a list of cells for each trajectory and their related pseudotime values were extracted from Monocle3 and used as input for tradeSeq. The 4,387 genes identified were further clustered into 24 modules (m1-m24) of co-expression along the eight major cell trajectories using the find_gene_modules function from the Monocle3 R package.

### RNAscope *in situ* hybridization

The spatial distribution of cell subpopulations was investigated by *in situ* hybridization using the RNAscope® technology. Freshly collected human fetal gonads were fixed in 4% paraformaldehyde at 4°C for 24 hours, embedded in paraffin, and sectioned to a thickness of 5 μm. *In situ* hybridization was next performed using the RNAscope® Multiplex Fluorescent v2 assay. Briefly, sections were pretreated with H_2_0_2_ for 10 min at room temperature, boiled in 1X Target Retrieval buffer for 15 min and submitted to a Protease Plus treatment for 30 min at 40°C. The following steps, i.e. hybridization of probes, amplification and signal development, were performed according to the manufacturer’s instructions. Signals were detected using three different fluorophores, Opal 520 (1:500), Opal 620 (1:750) and Opal 690 (1:750) (Akoya Biosciences) diluted in TSA buffer. Finally, slides were covered with ProLong™ Gold Antifade Mountant with DAPI (Thermo Fisher Scientific), images were acquired on a ZEISS LSM 880 with Airyscan confocal laser microscope (Zeiss) and processed using the ZEN 2.3 lite software (Zeiss).

### Tissue clearing and light-sheet imaging

After collection, samples were fixed overnight in 4% paraformaldehyde at 4°C, priori to being rinsed and stored in 1X PBS at 4°C. Samples were then dehydrated at room temperature in a graded series of MeOH concentrations in 1X PBS (20%, 40%, 60%, 80% and 2x 100%) with agitation before bleaching by overnight incubation in 6% H_2_O_2_ in 100% MeOH at 4°C shielded from light. Samples were washed 3 times in 100% MeOH before being rehydrated in a decreasing concentration of MeOH in 1X PBS (80%, 60%, 40%, 20%) with agitation before blocking and permeabilization in PBSGT (1X PBS with 0.2% gelatin and 0.5% Triton X-100) overnight. Following blocking, samples were incubated with primary antibodies against PAX8 (Proteintech, 10336-1-AP) and SOX9 (R+D Systems, AF3075) or Oct3/4 (R+D Systems, AF1759) at 1:500 dilution in PBSGT with Saponin (10mg/mL) with agitation for 7 days. Samples were then washed 6 times in PBSGT before incubation with secondary antibodies (ab150063 and ab175745 from Abcam, at 1:500 and 1:250 dilution, respectively) diluted in PBSGT with Saponin at 37°C with agitation for 2 days. After washing 6 times in PBSGT samples were embedded in 1.5% Agarose in 1X TAE buffer to facilitate mounting. Embedded samples were then dehydrated in a graded series of MeOH concentrations in 1X PBS (20%, 40%, 60%, 80% and 2 times 100%) with rotation before overnight incubation in 2/3 Dichloromethane (DCM) followed by a 30-minute incubation in 100% DCM before immersion in Benzyl ether (DBE) and stored at RT in the dark until imaging. Light-sheet imaging was performed on an Ultramicroscope Blaze light-sheet (Miltenyi Biotech) equipped with a 12x objective (MI Plan 12×/0.53) and tile scans of the whole samples were performed using a z-step size of 2µm.

### Data availability

Sequencing data and the gene count matrix were deposited at the European Genome-phenome Archive (EGA) (Freeberg *et al*, 2022) under the accession number EGAS00001006568. All processed data are also available through the ReproGenomics Viewer (RGV; https://rgv.genouest.org) database (Darde *et al*, 2015, 2019).

## Author contributions

ADR and FC supervised the research. AL, ADR, FC, and SMG designed the study. AL, FC and ADR wrote the manuscript. AL, ADR and FC analyzed and interpreted sequencing data. SL and OC contributed to the analysis. AL, ADR, SMG, BE and IC prepared the biological samples and interpreted sequencing data. AS and SMG validated expression data. CLM and AC performed tissue clearing and light-sheet imaging. SMG, SN and AC contributed to the manuscript. All authors approved the final version of the manuscript.

## Acknowledgements

Dedicated to the memory of our dear colleague, mentor and friend, Bernard Jégou. We thank all the staff of the Department of Obstetrics and Gynecology of the Rennes Sud Hospital for their expert assistance and help, and the participating women, without whom this study would not have been possible. We thank the GenOuest bioinformatics facility for hosting the softwares as well as all members of the Research Institute for Environmental and Occupational Health (IRSET) for stimulating discussions. We also thank the Institut national de la santé et de la recherche médicale (Inserm), the Université de Rennes 1, and the French School of Public Health (EHESP) for supporting this work. We thank S. Dutertre X. Pinson at the Microscopy Rennes Imaging Center (MRic, Biologie, Santé, Innovation Technologique -BIOSIT, Rennes, France) for assistance. MRic is a member of the national infrastructure France-BioImaging supported by the French National Research Agency (ANR-10-INBS-04). Part of the sequencing was performed by the GenomEast platform, a member of the ‘France Génomique’ consortium (ANR-10-INBS-0009).

## Contact for reagent and resource sharing

Requests for further information and resources may be directed to Antoine D. Rolland and Frédéric Chalmel.

## Supplementary information

### Key resources table

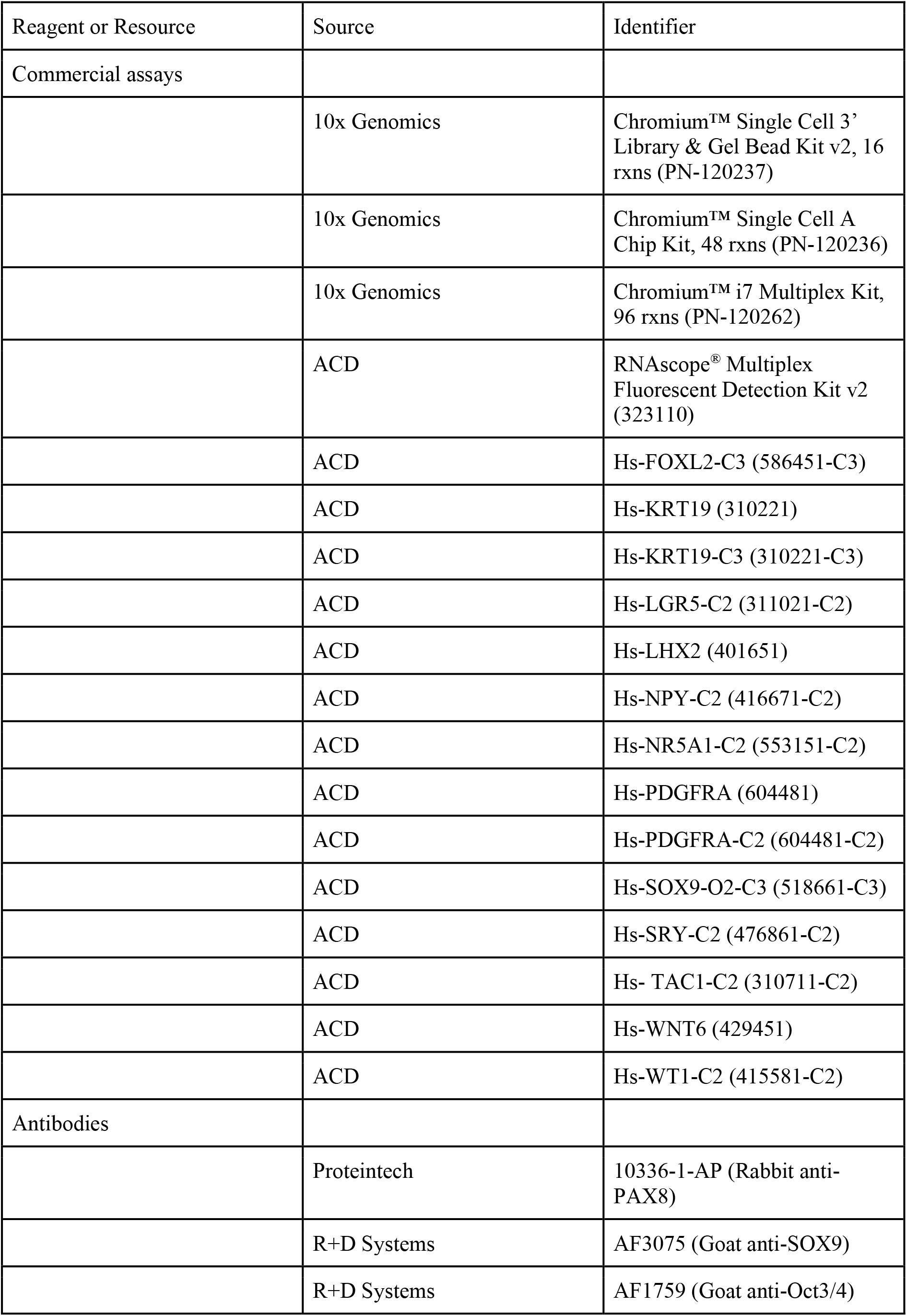
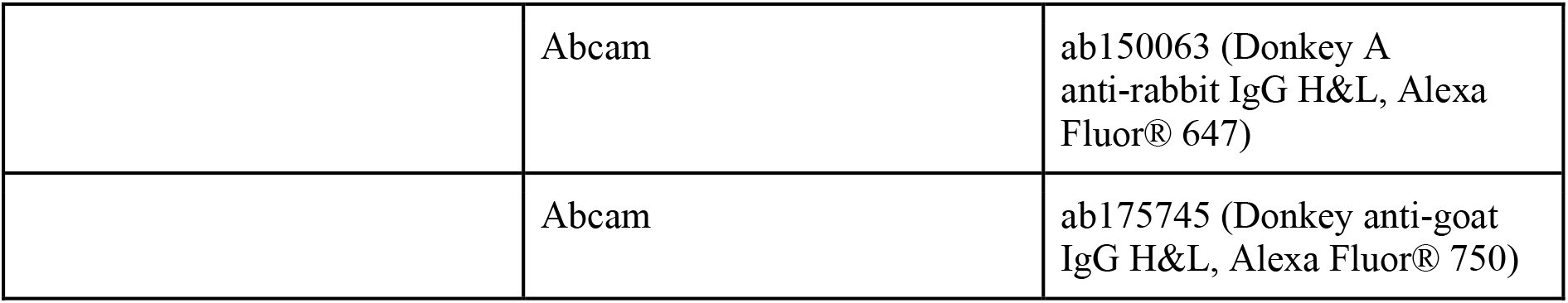

### Supplementary results

#### Section 1: Quality control and data preprocessing

We characterized the single-transcriptome of 33 fetal gonads (ranging from 5 to 12 PCW) by using the 10x Genomics single-cell RNA sequencing technology. After library preparation and sequencing, our initial dataset consisted of a total of 158,663 cells with an average of 77,769 reads per cell and a median number of 3,762 detected genes per cell. After quality control filtering, our final dataset described the expression of 29,323 genes in 127,903 single cells, with an average of 3,876 cells per and of 3,935 detected genes per cell (Table S1).

#### Section 2: Gonadal cell cluster annotation

To facilitate their annotation, we characterized each cell cluster i) in terms of gender and developmental stage (Fig. 2, panel C), ii) by assigning the cycle phase they belong to (G1, S or G2/M) (Fig. 2, panel C) and, iii) by evaluating the expression of known marker genes (Fig. 2, panels B and D; Table S2). In agreement with the embryonic primitive to definitive haemoglobin switch occuring during fetal life in humans at this developmental time frame (Sankaran *et al*, 2010), we identified two subpopulations of erythrocytes (Fig. 2, panels A, B and C). These included embryonic primitive erythrocytes (cluster c32), that are mostly found (>78%) in early developmental stages (5-6 PCW) and characterized by the expression of *HBE1*, as well as definitive red blood cells (cluster c39), that express *HBB* and are found in gonads from 8 PCW onwards. We also identified cluster c35 as endothelial cells, with enriched expression of angiogenic marker genes (*CD34* and *ECSCR*) whereas cluster c26 was significantly associated with genes related to immune cells (*PTPRC* and *TYROBP*) (Fig. 2, panel B). Cluster c41 appeared to correspond to perivascular cells, as evidenced by a very high expression of *TAGLN* and *ACTA2* (Fig. 2, panel B).

Three clusters (c33, c5 and c38) were identified as germ cells in agreement with the expression of several known markers for primordial germ cells (*POU5F1* and *NANOG*), gonadal germ cells (*DAZL*) and meiotic germ cells (*SYCP3*) (Fig. 2, panel B). The majority (>79%) of germ cells originated from female samples. Even though germ cells were present in all samples from as early as 5 PCW, cluster c5 was mainly composed of cells from mid-developmental stages (8-10 PCW) whereas cells in cluster c38 almost exclusively originated from ovaries at late developmental stages (10-12 PCW) (Fig. 2, panel C). Furthermore, many clusters could be associated with different gonadal somatic cell types based on the expression of markers, on top of genes known to be essential for gonad formation such as the transcription factors *GATA4*, *NR5A1*, *LHX9* and *WT1* (Lin & Capel, 2015). For instance, clusters c18, c29 and c9 were identified as coelomic epithelial cells expressing known markers such as *KRT19* and *UPK3B* and clusters c22 and c10 were identified as ovarian surface epithelial cells expressing *LHX2* and *IRX3* on top of *KRT19* and *UPK3B* (Mazaud *et al*, 2002; Tang *et al*, 2008; DeFalco *et al*, 2011; Sasaki *et al*, 2021; Kanamori-Katayama *et al*, 2011) (Fig. 2, panels B and D). While more than 85% of cells in clusters c9 and c18 were present in early developmental stages (5-6 PCW), c29 was predominantly associated with mid and late developmental stages (>72% in 8-12 PCW). In agreement with the high proliferation rate of coelomic epithelial cells as they give rise to the gonadal primordium (Yang *et al*, 2018), cluster c9 contained about 77% of cells in S or G2/M phases (Fig. 2, panel C). Additionally, several clusters of gonadal interstitial cells expressing high level of mesenchymal markers (*PDGFRA* and *TCF21*) could also be distinguished, including c1, c4, c13, c15, c16, c21, c23 and c25 (Fig. 2, panels A and B). Cluster c25 included cells from XX and XY gonads at all developmental stages, while cells of clusters c13 and c21 mostly originated from early developmental stages (6-7 PCW), with c21 and c25 corresponding to highly mitotically active interstitial cells (S or G2/M). Cluster c15 contained cells at early male development stages (6-7 PCW), while clusters c1, c16 and c23 were restricted to male gonads from mid and late developmental stages (7-12 PCW) and most probably correspond to interstitial precursors that will further differentiate into fetal Leydig cells. We actually identified two subpopulations of fetal Leydig cells, clusters c27 and c34, expressing higher levels of *INSL3* and of steroidogenic factors such as *CYP17A1*, respectively. Among interstitial cell clusters, only c4 was specific to female samples and is likely to include precursors of theca cells that will arise much later during ovarian development. As expected and unlike in male gonads, female interstitial cells therefore did not yet express canonical steroidogenic markers (Fig. 2, panel B).

Similarly, several clusters could be associated with supporting cell subpopulations (i.e. c2, c6, c7, c8, c11, c12, c14, c20, c24 and c31), based on the expression of *NR5A1* and *GATA4* and on the absence of mesenchymal markers. With the exception of c2 that included XX as well as XY gonadal cells, all clusters were restricted to either female (c6, c7, c8, c11 and c24) or male (c12, c14, c20 and c31) gonads. Clusters c2 and c24, mainly originating from 6 PCW and 7 PCW respectively, expressed *DMRT1 (Gierl et al, 2012; Warr et al, 2012)* and could be identified as early supporting cells, prior to the expression of the male or female canonical markers such as *SRY*, *SOX9* and *FOXL2*.Cluster c12 was characterized by peak expression of *SRY*, therefore representing pre-Sertoli cells, while clusters c14, c20 and c31 corresponded to more differentiated Sertoli cells that express higher levels of *SOX9* and *AMH* (Fig. 2, panel B). The canonical pre-granulosa cell marker *FOXL2* was expressed together with *IRX3* (Jorgensen & Gao, 2005; Fu *et al*, 2018) from 7 PCW onwards in clusters c6, c7, c8 and c11 (Fig. 2, panels B and C). Strikingly, *LHX9* was expressed in gonadal coelomic epithelial cells, ovarian surface epithelial cells, pre-granulosa cells and all gonadal interstitial cells, but not in male supporting cells(Fig. 2, panels B and C).

The exact nature of the abundant cluster c3 (6824 cells) was intriguing, as these cells neither expressed supporting nor interstitial markers (Fig. 2, panel C) (Birk *et al*, 2000; Gierl *et al*, 2012). Considering these cells were almost exclusively present in early gonads at 5-6 PCW (∼94%), expressed *NR5A1*, *GATA4* and *LHX9* but not *UPK3B* or *KRT19*, together with their central position within the UMAP, i.e. in-between early interstitial (cluster c15) and early supporting (clusters c2 and c12) cells (Fig. 2, panels A,B and C), we speculated that cluster c3 represented progenitors of gonadal somatic cell lineages. We considered clusters c19, c28 and c36 as mesonephric mesenchymal (*TCF21*+, *PDGFRA*+, *GATA2*+, *GATA4*-, *NR5A1*-, *LHX9*-), non-gonadal coelomic epithelial (*UPK3B*+, *KRT19*+, *GATA2*+, GATA4-, NR5A1-, LHX9-) and adrenocortical cells (*NR2F1*+, *CPLX1*+, *NR5A1*+, *GATA4*-, *LHX9*-), respectively. Cluster c40 was identified as neuroendocrine cells as they expressed *GATA2*, *ASCL1* and *PHOX2B* but not *GATA4*, *NR5A1* or *LHX9*. By investigating the expression of known markers for ducts, we found that the small cluster c37 corresponded to genital duct cells, as these cells expressed *KRT19*, *LHX1* and *PAX8* but neither *GATA4* nor *NR5A1* (Fig. 2, panel B). The consistent cluster c17 (3128 cells) on the other hand expressed *NR5A1*, *GATA4*, *PAX8* and *TBX1*, but not *LHX1*. This molecular signature strongly suggests that cluster c17 actually correspond to *rete testis* and *ovarii* cells, as recently characterized in the mouse (Kulibin & Malolina, 2020), and was confirmed by 3D imaging of PAX8-expressing cells using light sheet fluorescence microscopy on cleared whole-gonads (Fig. 3). In the end, only cluster c30 could not be associated with any known gene marker and corresponding cell type. As suggested by the low number of UMI counts in these cells, this cluster c30 was likely to correspond to residual low quality cells that tend to aggregate together within the UMAP.

### Supplementary figures

**Figure S1.**
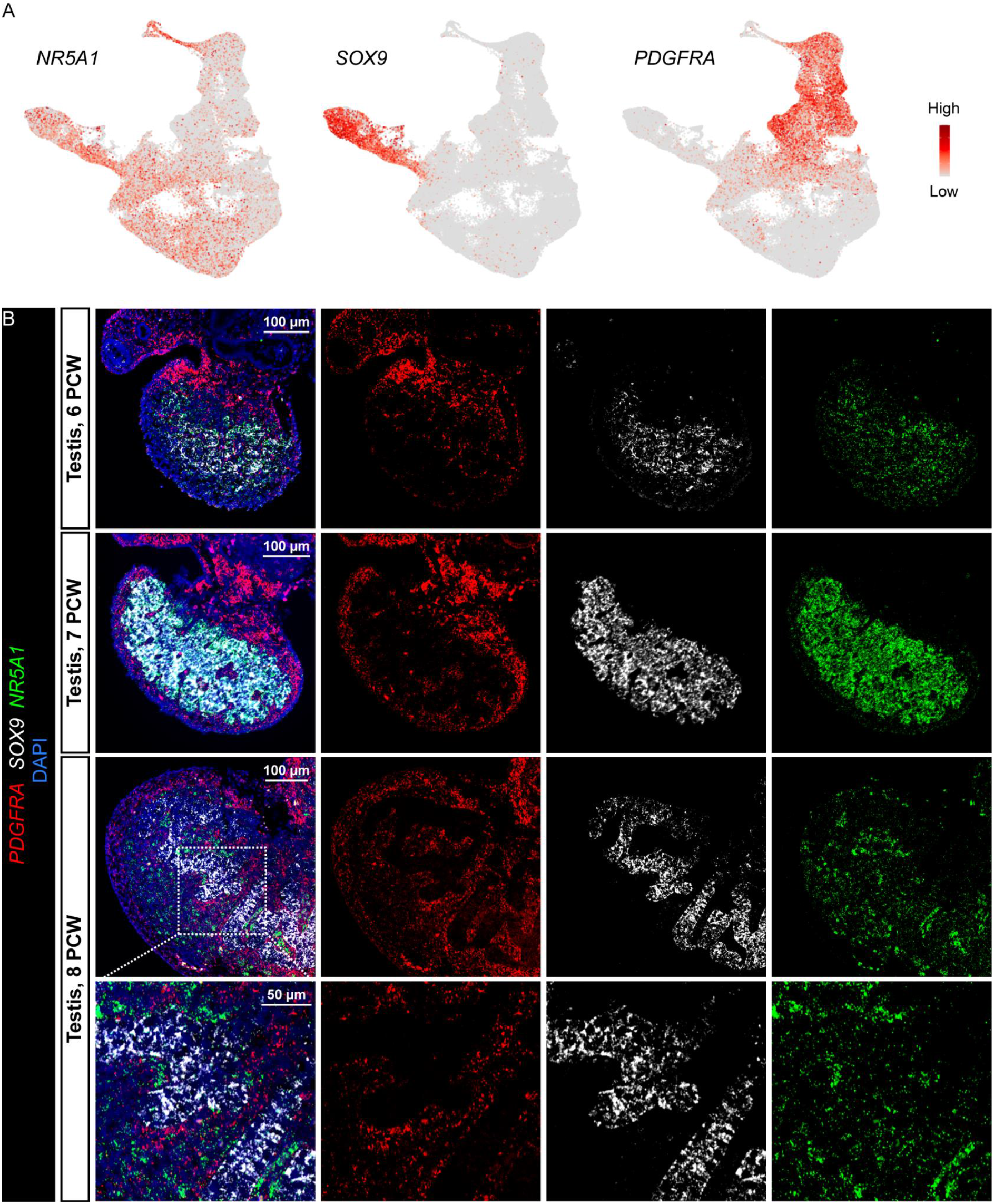
Expression dynamics of *NR5A1* during early human fetal testis development. **A.** UMAP visualization of the expression of *NR5A1* (Gonadal somatic cells), *SOX9* (Sertoli cells) and *PDGFRA* (Interstitial cells). **B.** RNAScope *in situ* hybridizations for *PDGFRA* (red), *SOX9* (white) and *NR5A1* (green) on sections from 6-8 PCW testes.

**Figure S2.**
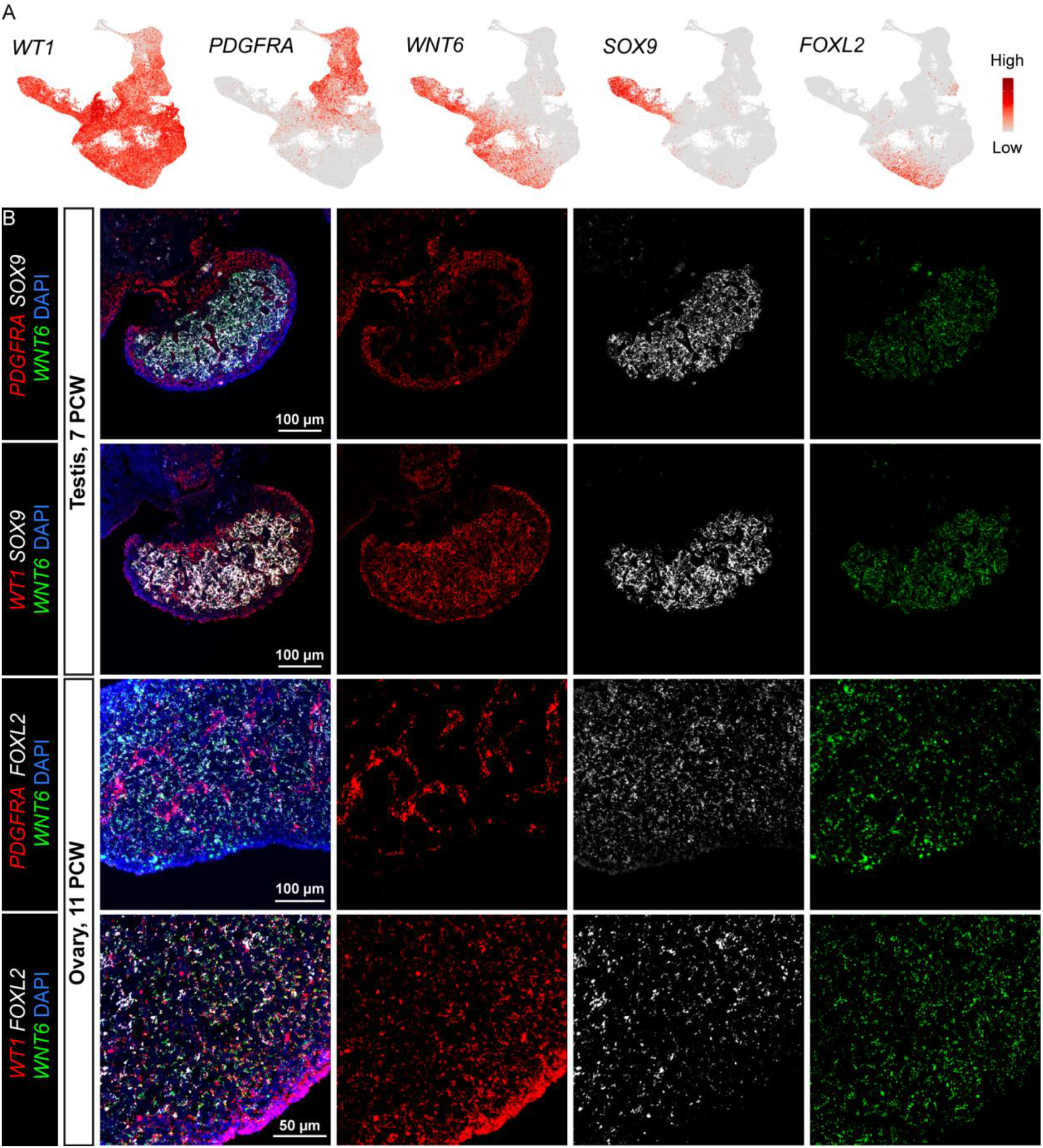
*WNT6* expression in male and female supporting cells of human fetal gonads. **A.** UMAP visualization of the expression of *WT1* (Gonadal somatic cells, except Leydig cells), *PDGFRA* (Interstitial cells), *WNT6* (Supporting cells), *SOX9* (Sertoli cells) and *FOXL2* (Pre-granulosa cells). **B.** RNAscope *in situ* hybridizations for *PDGFRA* (red) or *WT1* (red), *SOX9* (white) or *FOXL2* (white) and *WNT6* (green) on sections from a testis and an ovary at 7 and 11 PCW, respectively.

**Figure S3.**
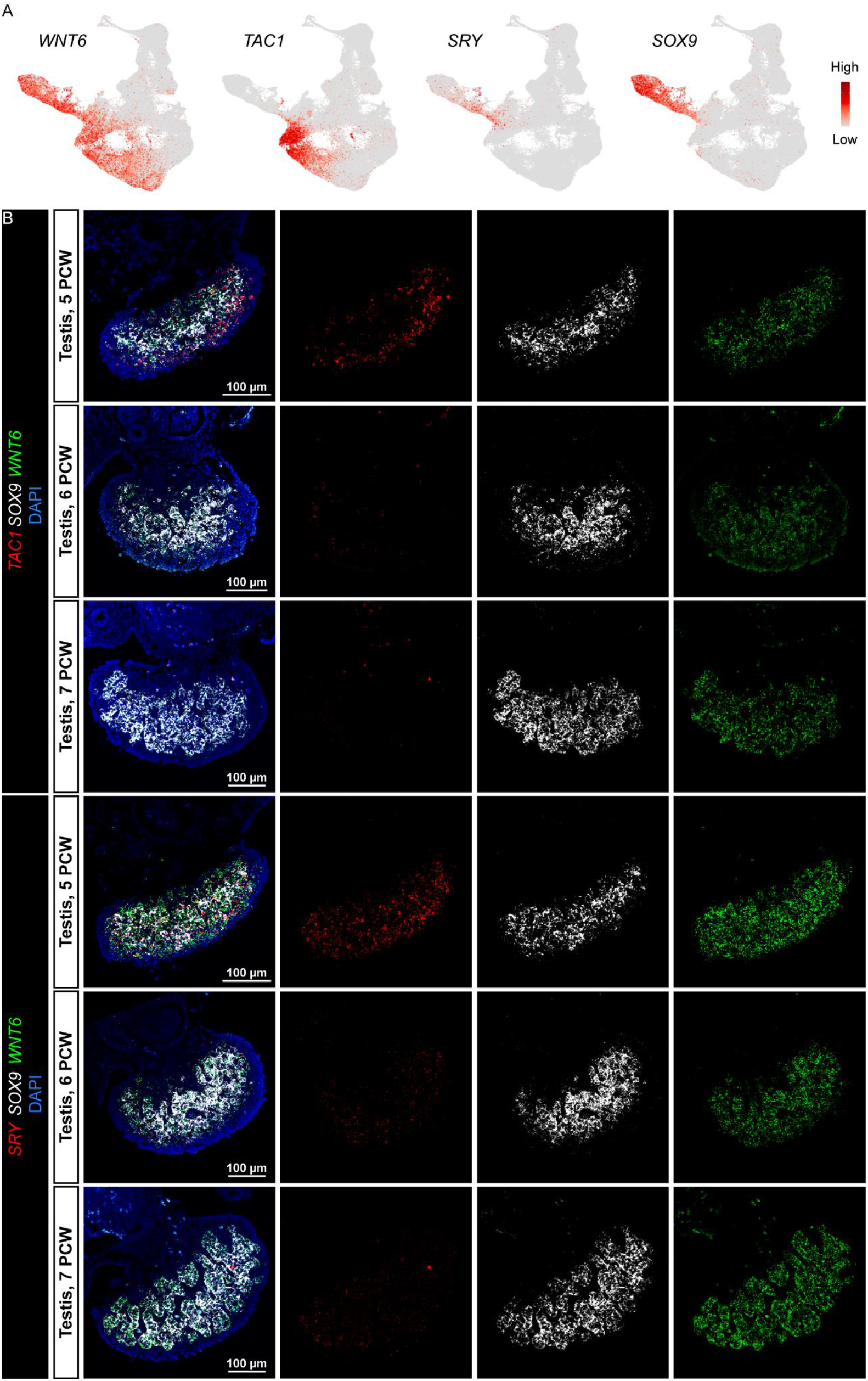
Transient expression of *TAC1* in early male supporting cells. **A.** UMAP visualization of the expression of *WNT6* (Supporting cells), *TAC1* (Early supporting cells), *SRY* (Pre-Sertoli cells) and *SOX9* (Sertoli cells). **B.** RNAscope *in situ* hybridizations for *SRY* (red) or *TAC1* (red), *SOX9* (white) and *WNT6* (green) on sections from human fetal testes at 5, 6 and 7 PCW.

**Figure S4.**
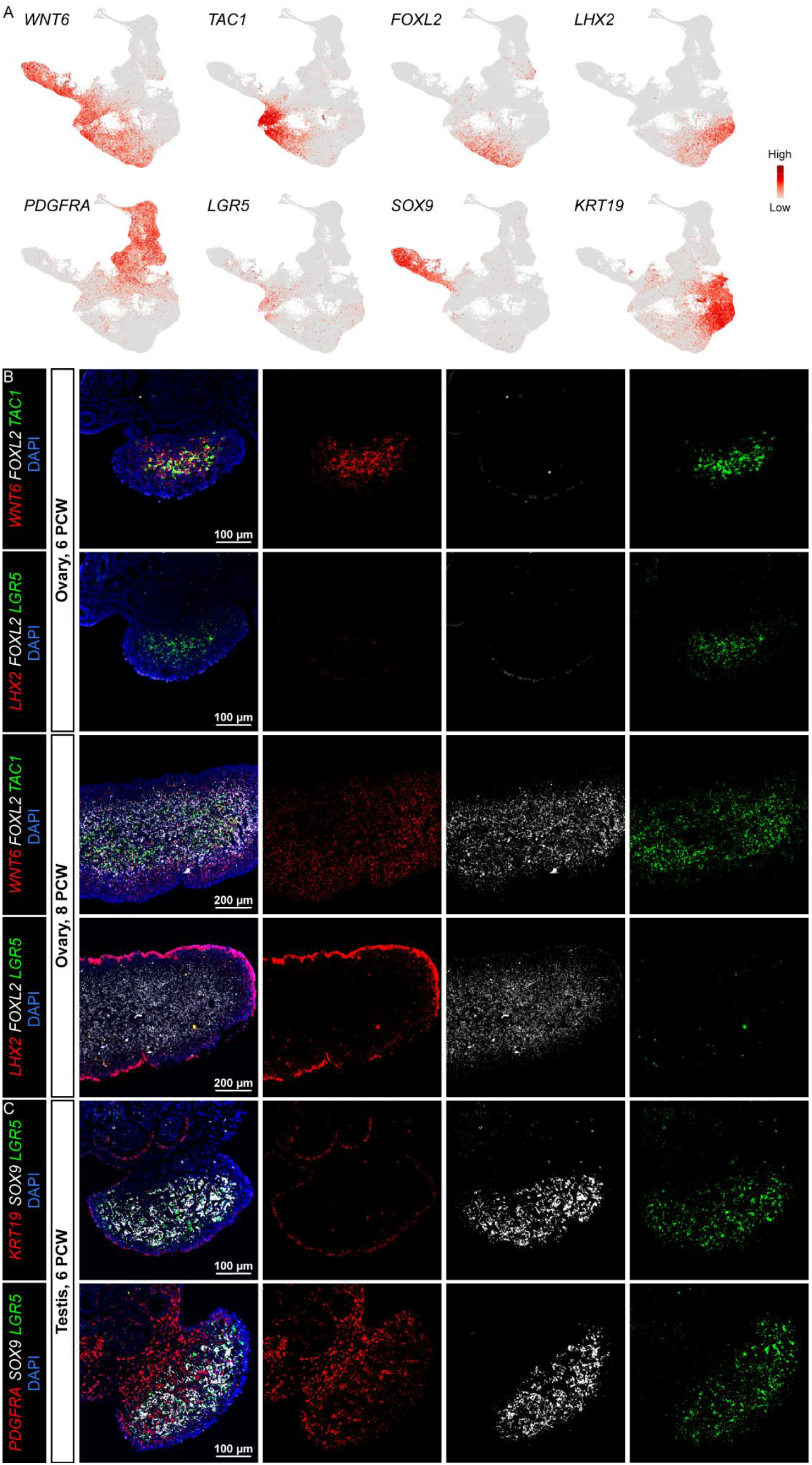
Expression of *LGR5* in early supporting cells of human fetal gonads. **A.** UMAP visualization of the expression of *WNT6* (Supporting cells), *PDGFRA* (Interstitial cells), *TAC1* and *LGR5* (Early supporting cells), *FOXL2* (Pre-granulosa cells), *SOX9* (Sertoli cells), *LHX2* (Ovarian surface epithelial cells and pre-granulosa cells) and *KRT19* (Coelomic epithelial cells and ovarian surface epithelial cells). **B.** RNAscope in situ hybridizations for *WNT6* or *LHX2* (red), *FOXL2* (white) and *TAC1* or *LGR5* (green) on sections from ovaries at 6 and 8 PCW. **C.** RNAscope in situ hybridizations for *KRT19* or *PDGFRA* (red), *SOX9* (white) and *LGR5* (green) on sections from a testis at 6 PCW.

**Figure S5.**
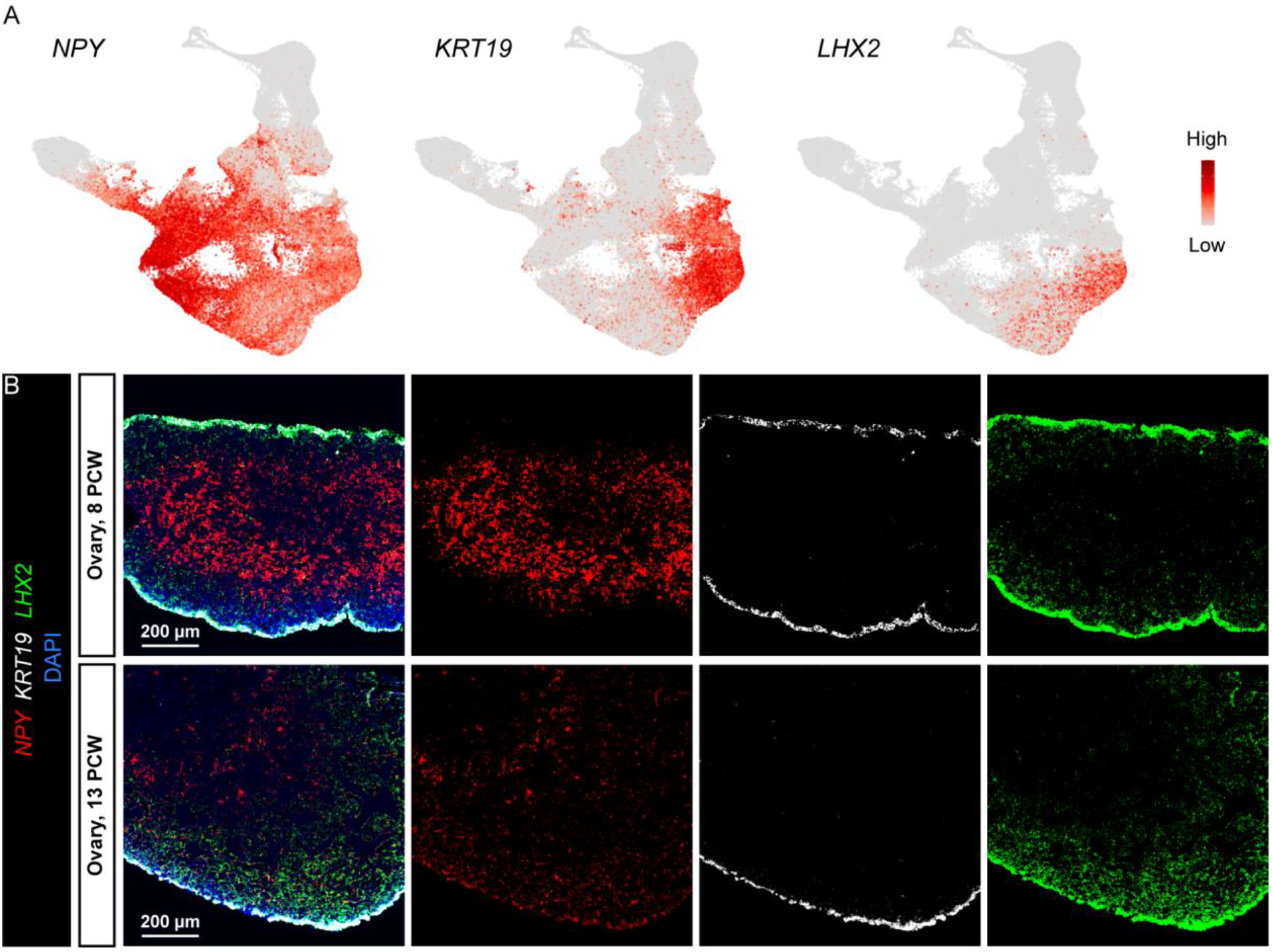
*LHX2* versus *KRT19* expression in human fetal ovaries. **A.** UMAP visualization of the expression of *NPY* (Coelomic epithelial cells, ovarian surface epithelial cells, gonadal progenitor cells, ovarian supporting cells, pre-Sertoli cells, *rete testis* and *rete ovarii* cells), *KRT19* (Coelomic epithelial cells and ovarian surface epithelial cells) and *LHX2* (Ovarian surface epithelial cells and pre-granulosa cells). **B.** RNAscope in situ hybridizations for *NPY* (red), *KRT19* (white) and *LHX2* (green) on sections from ovaries at 8 and 13 PCW.

### Supplementary table legends

**Table S1. Description of single-cell RNA-seq samples.**

Development stage in PCW, gender and sequencing data are presented for each sample.

**Table S2. Differentially expressed genes (DEGs) between the 41 clusters (c1-c41) in the total dataset.**

DEGs were identified using the Wilcox rank sum test implemented in Seurat package (v4.0.1) (one-tailed Wilcoxon rank sum test, p-values adjusted for multiple testing using the Bonferroni correction).

**Table S3. Differentially expressed genes (DEGs) between the 42 somatic cell clusters (s1-s42) in the dataset of somatic cells.**

DEGs were identified using the Wilcox rank sum test implemented in Seurat package (v4.0.1) (one-tailed Wilcoxon rank sum test, p-values adjusted for multiple testing using the Bonferroni correction).

**Table S4. Differentially expressed genes (DEGs) over the 8 major cell trajectories.**

DEGs over the 8 major pseudotemporal cell trajectories were determined with the tradeSeq package with a number of knots (nknots) = 6.

